# The structural repertoire of *Fusarium oxysporum* f. sp. *lycopersici* effectors revealed by experimental and computational studies

**DOI:** 10.1101/2021.12.14.472499

**Authors:** Daniel S. Yu, Megan A. Outram, Ashley Smith, Carl L. McCombe, Pravin B. Khambalkar, Sharmin A. Rima, Xizhe Sun, Lisong Ma, Daniel J. Ericsson, David A. Jones, Simon J. Williams

**Author notes:** Corresponding author: Simon Williams, Megan Outram. Current address: Black Mountain Science and Innovation Park, CSIRO Agriculture and Food, Canberra, Australia (Megan A. Outram). Funding: Australian Research Council (DP200100388, FT200100135, DE170101165). Australian Academy of Science (Thomas Davies Grant). Australian National University Future Scheme (35665). Australian Institute of Nuclear Science and Engineering.

## Abstract

Plant pathogens secrete proteins, known as effectors, that function in the apoplast or inside plant cells to promote virulence. Effector recognition by cell-surface or cytosolic receptors results in the activation of defence pathways and plant immunity. Despite their importance, our general understanding of fungal effector function and recognition by immunity receptors remains poor. One complication often associated with effectors is their high sequence diversity and lack of identifiable sequence motifs precluding prediction of structure or function. In recent years, several studies have demonstrated that fungal effectors can be grouped into structural classes, despite significant sequence variation and existence across taxonomic groups. Using protein x-ray crystallography, we identify a new structural class of effectors hidden within the secreted in xylem (SIX) effectors from *Fusarium oxysporum* f. sp. *lycopersici* (*Fol*). The recognised effectors Avr1 (SIX4) and Avr3 (SIX1) represent the founding members of the ***Fol* d**ual-domain (FOLD) effector class, with members containing two distinct domains. Using AlphaFold2, we predicted the full SIX effector repertoire of *Fol* and show that SIX6 and SIX13 are also FOLD effectors, which we validated experimentally for SIX6. Based on structural prediction and comparisons, we show that FOLD effectors are present within three divisions of fungi and are expanded in pathogens and symbionts. Further structural comparisons demonstrate that *Fol* secretes effectors that adopt a limited number of structural folds during infection of tomato. This analysis also revealed a structural relationship between transcriptionally co-regulated effector pairs. We make use of the Avr1 structure to understand its recognition by the I receptor, which leads to disease resistance in tomato. This study represents an important advance in our understanding of *Fol-*tomato, and by extension plant-fungal interactions, which will assist the development of novel control and engineering strategies to combat plant pathogens.

## Introduction

*Fusarium oxysporum* is a soil-borne fungal pathogen responsible for destructive vascular wilt diseases in a wide range of plants. It ranks within the top ten important fungal pathogens in terms of scientific and economic importance [1]. The best-characterised *F. oxysporum* pathosystem involves *F. oxysporum* f. sp. *lycopersici* (*Fol*) and tomato. Previous studies of *Fol-*infected tomato identified a number of fungal proteins within the xylem sap [2]. These secreted in xylem (SIX) effector proteins represent major pathogenicity determinants across different *formae speciales* of *F. oxysporum*. Currently, 14 SIX effectors have been identified in *Fol* consisting of small (less than 300 amino acids in length), secreted, cysteine-rich proteins [3-6]. Most SIX effectors are encoded on the conditionally-dispensable chromosome 14 required for *Fol* pathogenicity [7]. This dispensable chromosome can be horizontally transferred from *Fol* to a non-pathogenic strain of *F. oxysporum*, resulting in a transfer of pathogenicity [4, 7]. To date, all 14 SIX effectors lack sequence identity with proteins of known function, preventing prediction of function based on their amino acid sequence. Several SIX effectors have been shown to be essential for full virulence including SIX1, SIX2, SIX3, SIX5 and SIX6 from *Fol* [5, 8-11], SIX1 from *F. oxysporum* f. sp. *conglutinans* (*Focn*), which infects cabbage [12], SIX4 from *F. oxysporum* isolate Fo5176, which infects Arabidopsis [13], and SIX1 and SIX8 from *F. oxysporum* f. sp. *cubense*, which infects banana [14, 15]. *Fol SIX3* (*Avr2*) and *SIX5* are adjacent, divergently-transcribed genes with a common promoter, and SIX5 has been shown to interact with SIX3 to promote virulence by enabling symplastic movement of SIX3 via plasmodesmata [16]. *Focn SIX8* and *PSE1* (pair with *SIX8* 1) are also a divergently-transcribed effector gene pair that function together to suppress phytoalexin production and plant immunity in Arabidopsis [17]. In *Fol, SIX8* forms a similar gene pair with *PSL1* (*PSE1*-like 1) [17]. Despite their roles in fungal pathogenicity, the virulence functions of most SIX effectors remain unknown.

To combat pathogen attack, plants possess resistance genes that encode immunity receptors capable of recognising specific effectors leading to disease resistance. Four resistance genes, introgressed into tomato from related wild species, have been cloned. *I* and *I-7* encode transmembrane receptor proteins containing extracellular leucine-rich repeat (LRR) domains and short cytoplasmic domains (LRR-RPs) [18, 19]. *I-2* encodes a cytoplasmic receptor containing nucleotide binding (NB) and C-terminal LRR domains [20], while *I-3* encodes a transmembrane protein with an extracellular S-receptor-like domain and cytoplasmic serine/threonine kinase domain (SRLK) [21]. *Fol* Avr1 (SIX4), Avr2 (SIX3) and Avr3 (SIX1) are recognised by tomato immunity receptors, I, I-2 and I-3, respectively, leading to effector-triggered immunity (ETI) and disease resistance [5, 22, 23].

By understanding the function of *F. oxysporum* effector proteins, and how specific effectors are detected by immunity receptors, we (and others) hope to develop novel disease management strategies targeting vascular wilt diseases. Protein structure studies of effectors provide one avenue to assist this pursuit. Currently, Avr2 represents the only SIX effector whose protein structure has been determined [24]. Interestingly, the β-sandwich fold of Avr2 revealed that this effector shares structural homology to ToxA from *Pyrenophora tritici-repentis* and AvrL567 from *Melampsora lini* [25, 26], despite a lack of sequence identity. The observation of structural classes for effectors without identifiable domains or homologies to proteins of known function has been demonstrated experimentally for four effector structural families, including the so-called MAX (***M****agnaporthe oryzae* **A**vr effectors and To**x**B from *P. tritici-repentis*) [27], RALPH (**R**N**A**se-**L**ike **P**roteins associated with **H**austoria) [28], LARS (***L****eptosphaeria* **A**vi**r**ulence and **S**upressing) [29] and ToxA-like families [24-26].

Combining experimental and computational approaches, we present the structural repertoire of sequence unrelated effectors utilised by *Fol* during infection of tomato, including the classification of a new effector family, the FOLD (***Fol* d**ual-domain) effectors. We show using structural comparisons that FOLD effectors are widely distributed in phytopathogenic fungi as well as symbionts. Further, we define the domains and residue that mediate the recognition of the FOLD effector, Avr1, by its corresponding immunity receptor.

## Results

### The structures of Avr1 and Avr3 adopt a similar dual-domain fold

Avr1 and Avr3 are cysteine-rich effectors that belong to the K2PP (Kex2-processed pro-domain) effector class [30, 31]. To help understand their function, and recognition by I and I-3, we sought to solve their structures using x-ray crystallography. Using our optimised protein production strategy [32], we produced Avr1 (Avr1^18-242^) and Avr3 (Avr3^22-284^) in *Escherichia coli* for crystallisation studies (S1A and S1B Fig). Crystals were obtained for Avr3^22-284^ (after referred to as Avr3) (S1B Fig), however, Avr1^18-242^ failed to crystallise. Previously, we demonstrated that pro-domain removal from the K2PP effector SnTox3 was required to obtain protein crystals [30] and predicted this may also be important for Avr1. Treatment of Avr1^18-242^ with Kex2 *in vitro* resulted in a predominant Avr1 band of ∼20 kDa consistent with a mature Avr1^59-242^ protein, however, lower molecular weight bands were also observed suggesting *in vitro* Kex2 cleavage at additional sites [30]. To address this, Avr1 was engineered with an internal thrombin cleavage site (replacing the Kex2 site) to produce a single Avr1^59-242^ product after thrombin cleavage (after referred to as Avr1). This protein was subsequently used for crystallisation studies resulting in rectangular plate-like crystals (S1A Fig).

The crystal structures of Avr1 and Avr3 were solved using a bromide-ion-based single-wavelength anomalous diffraction (SAD) approach (S1 Table), and subsequently were refined using a native dataset to a resolution of 1.65 Å and 1.68 Å, respectively (Fig 1A and 1B). Despite sharing low amino-acid sequence identity (19.5%), Avr1 and Avr3 adopt a structurally similar dual-domain protein fold. Interpretable, continuous electron density was observed from residue 96 in Avr3 and some regions of the intact pro-domain could be interpreted in the electron density (residues 26-49) (S2A Fig). We also identified regions of the pro-domain (residues 23-45) of Avr1 in the electron density, despite thrombin cleavage of the pro-domain prior to crystallisation (S1A Fig). This indicates that an association between respective Avr and pro-domain was maintained post cleavage *in vitro* (S2B Fig). The importance of this association, if any, remains unclear, but for simplicity, the pro-domains were excluded from further analysis.

**Fig 1.**
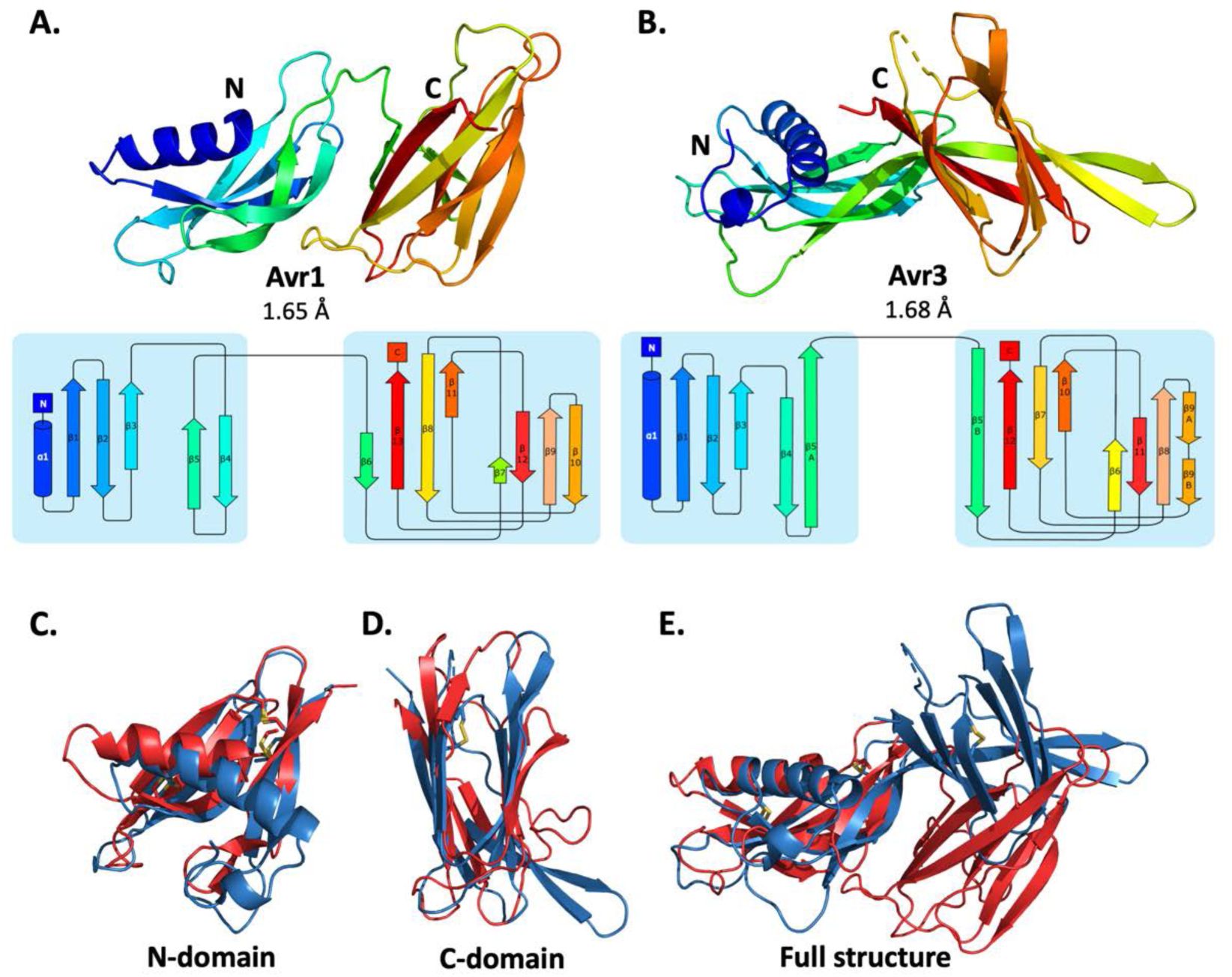
Crystal structures of Avr1 and Avr3 from *Fol* adopt a similar structural fold. Ribbon diagrams of **(A)** Avr1 and **(B)** Avr3 coloured from N- (blue) to C-terminus (red) showing the dual-domain structural fold (top panels), and secondary structure topology map (bottom panels) of Avr1 and Avr3, respectively. For both, the N-domain is shown on the left and the C-domain is shown on the right. The colours of the secondary structural elements match the colours depicted on the crystal structure. Structural alignments of Avr1 (shown in red) and Avr3 (shown in blue) showing **(C)** N-domains alone, **(D)** C-domains alone and **(E)** full structures. Disulfide bonds are shown in yellow. Structural alignment was performed using the pairwise alignment function of the DALI server [33].

The Avr1 and Avr3 N-terminal domain (N-domain), consisting of an N-terminal α-helix followed by five β-strands, and C-terminal domain (C-domain), consisting of a β-sandwich architecture, involving seven or eight β-strands are very similar with a root-mean-square deviation (RMSD) of 2.1 Å and 2.8 Å, respectively (superposition performed using DALI server [33]) (Fig 1). While the individual domains are very similar, superposition of the dual-domain structures returns an RMSD of ∼3.4 Å. The larger difference is due to a rotation between the N-and C-domains (Fig 1E). The structures of Avr1 and Avr3, when compared with the solved structures of other fungal effectors, demonstrate that they adopt a unique two-domain fold and represent the founding members of a new structural class of fungal effectors we have designated the FOLD (***Fol* d**ual-domain) effectors.

### SIX6 and SIX13 belong to the FOLD effector family

We were interested to determine if other SIX effectors belonged to the FOLD effector family. One conserved sequence feature observed in Avr1 and Avr3 was the spacing of the six cysteines within the N-domain. We analysed the cysteine spacing of the other SIX effectors and found that SIX6 and SIX13 contained a cysteine profile like Avr1 and Avr3 (Fig 2A), suggesting they may be FOLD effectors. With the recent advances in *ab initio* structural prediction by Google DeepMind’s AlphaFold2 [34] we predicted the structures of the SIX effectors to determine if, as suggested by our sequence analysis, other SIX effectors are FOLD effector family members.

**Fig 2.**
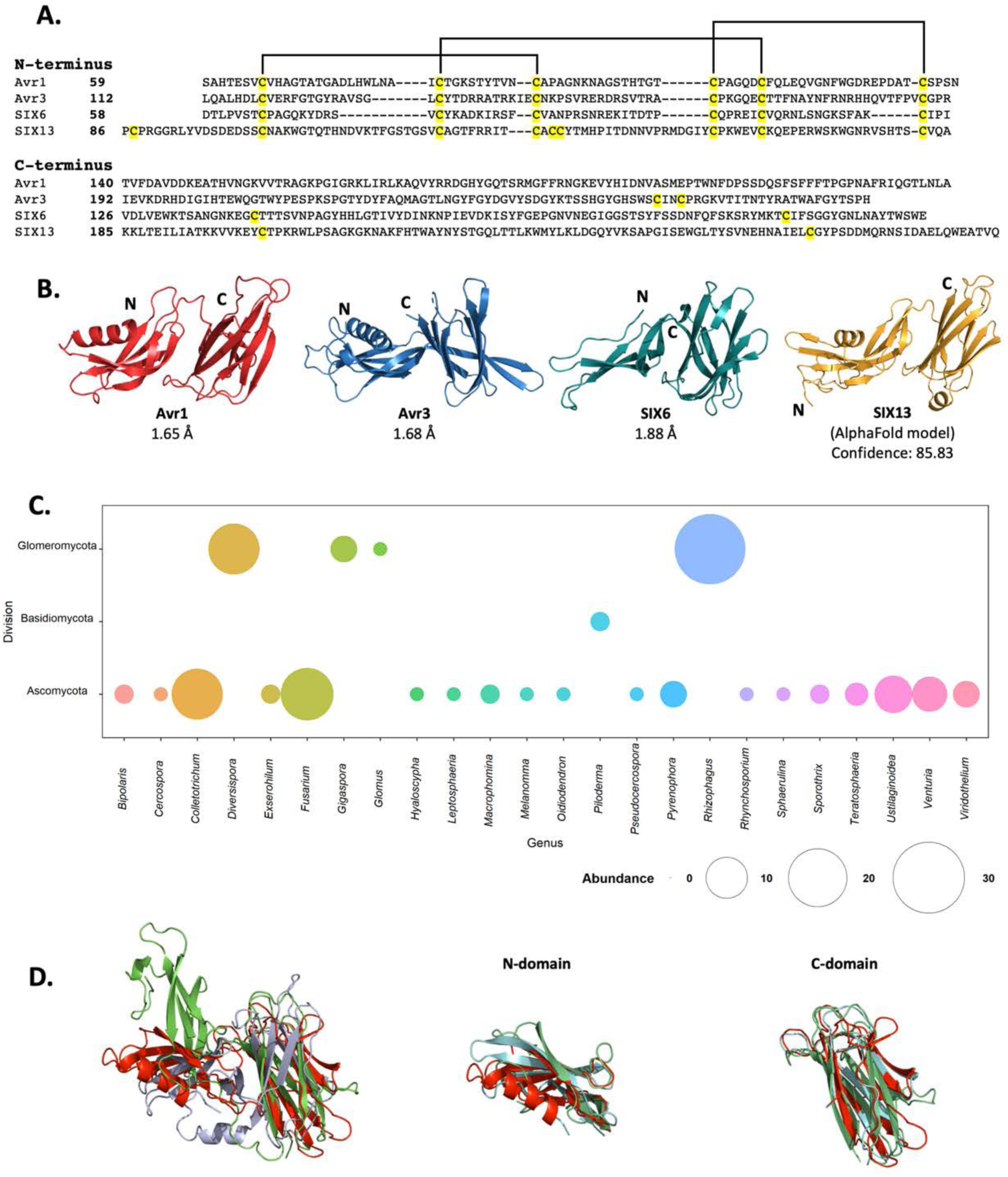
FOLD effector family is distributed within *Fusarium oxysporum* and other fungi. **(A)** Amino acid sequence alignment of the mature Avr1, Avr3, SIX6 and SIX13 sequences show a common cysteine spacing at the N-terminus. The alignment is split into the N-terminus (N-domain; top panel) and C-terminus (C-domain; bottom panel). Cysteine residues are highlighted in yellow, with common disulfide bonding connectivity, as determined by the crystal structures of Avr1 and Avr3, shown with black lines. **(B)** Ribbon diagrams of the Avr1, Avr3, SIX6 crystal structures and SIX13 AlphaFold2 model show a conserved dual-domain structure. The N-and C-termini are labelled. **(C)** Structure-guided search for putative FOLD effectors across fungi using Foldseek webserver. The size of the circles represent abundance with genus. **(D)** Superposition (structural alignment) of representative putative FOLD effectors from the divisions Glomeromycota and Basidiomycota with Avr1 in ribbon representation. Putative FOLD protein from *Rhizophagus clarus* (UniProt: A0A2Z6QDJ0) in light blue, and *Piloderma croceum* (UniProt: A0A0C3C2B2) in green. FOLD structural alignment (right), N-domain only (middle), C-domain only (right).

As an initial step we benchmarked the AlphaFold2 predicted models of Avr1 and Avr3 (downstream of the Kex2 cleavage site (Avr1^59-242^ and Avr3^96-284^) against our experimentally determined structures (S3 Fig). The AlphaFold2 model of Avr1 returned a low average per-residue confidence score (pLDDT = 55%) and the RMSD was 6.9 Å when model and structure were compared, however, the dual domain architecture was correctly predicted with a Z-score of 11.3 identified using a DALI pair-wise structural comparison (S3A Fig and S3E). The AlphaFold2 model of Avr3 returned a high average pLDDT score (92%) and superimposed well to the solved structure (S3B Fig), despite a slight skew between the orientation of the individual domains (RMSD = 3.6 Å overall; 1.1 Å for the N-domain; 0.8 Å for the C-domain). This demonstrated that accurate FOLD effector prediction was possible using AlphaFold2.

We subsequently generated SIX6 and SIX13 models, downstream of the predicted Kex2 cleavage site (SIX6^58-225^, SIX13^78-293^), using AlphaFold2 and obtained high average confidence scored models supporting their inclusion in the FOLD family (S4 Fig). To validate this experimentally, we produced SIX6^58-225^ and SIX13^22-293^ (after referred to as SIX6 and SIX13) as described for Avr1/Avr3 and obtained crystals for both proteins (S1 Fig). While the SIX13 crystals diffracted poorly, the SIX6 crystals diffracted x-rays to ∼1.9 Å and we solved the structure of SIX6 using the AlphaFold2 generated model as a template for molecular replacement (Fig 2B, S1 Table), confirming its inclusion as a member of the FOLD family. Despite lacking an N-terminal helix, the N-domain contains five β-strands held together by three disulfide bonds with an arrangement, identical to Avr1 and Avr3. The C-domain is an eight stranded β-sandwich that is stabilised by a single disulfide bond (unique to SIX6 compared to Avr1 and Avr3) connecting the β7 and β12 strands. Like Avr1, we identified regions of the pro-domain within the SIX6 structure (residues 29-46), despite cleavage of the pro-domain prior to crystallisation (S2C Fig), but only within one molecule in the asymmetric unit (S2D Fig). For structural analysis, we used the structured region of Chain A of SIX6 (Fig 2B).

### FOLD effectors are distributed across multiple fungal genera

Despite structural similarities, the FOLD effectors are divergent in their amino acid sequences, sharing 15.5 – 22.5% sequence identities between all members (Fig 2A). Homologues of FOLD effectors are dispersed across multiple *formae speciales* of *F. oxysporum* (S5 Fig) [6, 8, 35-38]. We were interested to understand the distribution of FOLD effectors in fungi. Previous structural-based searches performed on effector candidates from *Venturia inaequalis* using Avr1 and Avr3 as templates (which we provided to the authors) found three candidates predicted to be FOLD effectors [39]. Here, we utilised our experimentally determined structures (Avr1, Avr3 and SIX6) to search for other fungal FOLD effectors within the AlphaFold2 protein structure database [40] (https://alphafold.ebi.ac.uk/) using the Foldseek webserver [41]. This analysis identified 124 putative FOLD protein family members across three Divisions of Fungi (Ascomycota, Basidiomycota, and Glomeromycota) (Fig. 2C). Over half of these were found in Ascomycota fungi (73), with expanded families in species of *Fusarium* and *Colletotrichum* (Fig 2C, S2 Table). Expanded families of FOLD proteins were also observed in the division Glomeromycota that form arbuscular mycorrhiza in plant roots, while two putative FOLD effectors were also predicted in the ectomycorrhizal fungus *Piloderma olivaceum* (Division Basidiomycota), which forms mutualistic associations with conifer and hardwood species [41]. Structural superposition of members from the three Divisions confirms the structural similarities between the N and C domains and highlights that the major differences identified are the orientation of the domains relative to each other (Fig. 2D), consistent with our experimental data for Avr1, Avr3 and SIX6.

### Distinct structural families exist among the other SIX effectors

With the successful utilisation of AlphaFold2 as a model for molecular replacement (SIX6 structure), and structural similarity searches for FOLD effectors, we decided to perform structural comparisons with the remaining SIX effectors. AlphaFold2 modelling of the effectors was conducted on sequences with the signal peptide and putative pro-domain (if present) removed (S6 Fig). The models and experimentally determined SIX effector structures (Avr1, Avr2, Avr3 and SIX6) were compared using the DALI server [33] and a Z-score with a cutoff of >2 was used to indicate structure similarity.

The observed structural similarity between the FOLD effectors was high, with Z-scores above 8 for all comparisons (Fig 3A). Avr2, a member of the ToxA-like effector family, exhibited structural similarity with the SIX7 and SIX8 models (Z-scores > 5) (Fig 3A). Analysis of the models and topology show that SIX7 and SIX8 both consist of a β-sandwich fold, strongly indicating their inclusion of within the ToxA-like structural family (Fig 3C, S7 Fig).

**Fig 3.**
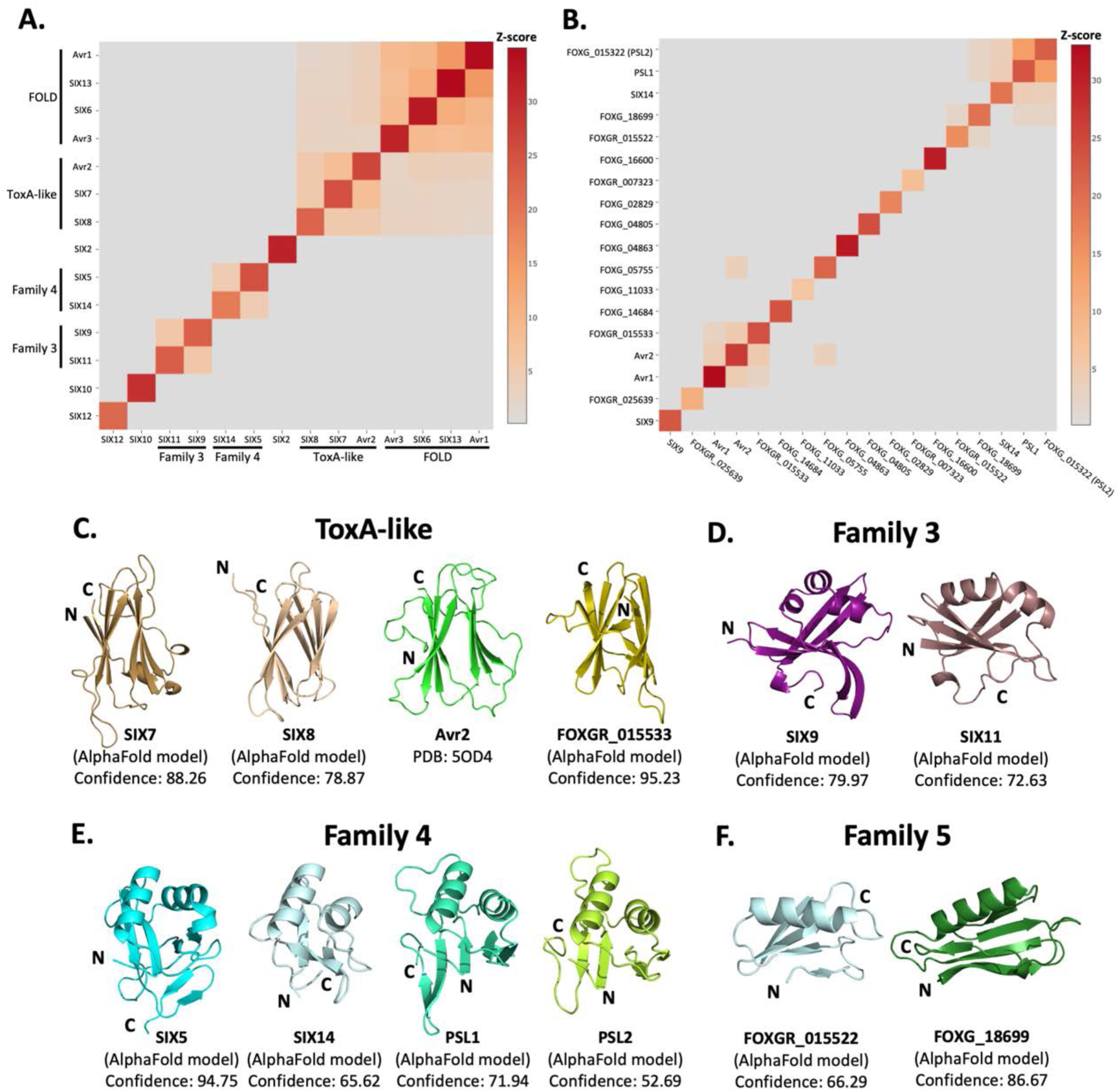
Identification of new putative structural families within the SIX effectors. Heat maps showing the structural similarity of structures and AlphaFold2 models of the **(A)** SIX effectors and **(B)** effector candidates from *Fol* in a structural pairwise alignment. Amino acid boundaries that were modelled for each protein is found in S3 Table. Structural similarity was measured with Z-scores. A cutoff Z-score of 2 was applied for defining structural families. Z-score scale is shown in a grey to red spectrum. **(C)** Cartoon representation of the ToxA-like effectors from *Fol*. AlphaFold2 models of SIX7, SIX8 and FOXGR_015533 effector candidate are putative members of the ToxA-like effector family. The crystal structure of Avr2 [24], another member of the ToxA-like effector family, is shown in green for comparison. Cartoon representations of **(D)** Family 3, **(E)** Family 4 and **(F)** Family 5 consisting of members that are predicted to be structurally similar. The N-and C-termini are labelled. Structural similarity searches were performed using the DALI server [33].

Beyond these described structural families, the Z-scores indicated that two additional, but not yet characterised, structural families exist within the SIX effectors. Here, we define these as structural family 3 and 4, consisting of SIX9 and SIX11, and SIX5 and SIX14, respectively (Fig 3D, E). The models of SIX9 and SIX11 both consist of five β-strands and either two or three α-helices (Fig 3D, S8 Fig), despite sharing only 14% sequence identity. To further our understanding of the putative function of this family we did a structural search against the protein databank (PDB) and found that both structures share structural similarity to various RNA binding proteins (Z-scores > 2.5), such as the RNA recognition motif (RRM) fold of the Musashi-1 RNA-binding domain (PDB code: 5X3Z) [42] with a Z-score of 2.6 and 4.5 for SIX9 and SIX11, respectively.

SIX5 and SIX14 also share limited sequence identity (23%) but the structural predictions show a similar secondary-structure topology consisting of two α-helices and four to six β-strands (Fig 3E, S8 Fig). We compared the models of SIX5 and SIX14 against the PDB using DALI and identified structural similarity toward the *Ustilago maydis* and *Zymoseptoria tritici* KP6 effector (PDB codes: 4GVB and 6QPK) [43], suggesting SIX5 and SIX14 belong to the KP6-like structural family (S7 Fig). Collectively, this analysis demonstrates that 11 of the 14 SIX effectors, group into 4 different structural families.

### Structural modelling and comparison of an expanded set of *Fol* effectors

The SIX effectors are only a subset of effectors utilised by *Fol* during infection of tomato. Recently, the *Fol* genome was re-sequenced [44] and reannotated in combination with RNAseq data from *Fol*- infected tomato plants [45]. A total of 26 genes encoding novel effector candidates were identified that were consistently upregulated during *Fol* infection [45], which were not previously predicted or predicted incorrectly in the original genome annotation [4]. Of these, 14 genes encoded proteins with no recognised domains or motifs based on their amino acid sequences. We generated structural models using AlphaFold2 of these 14 (S3 Table, S6 Fig) and structurally aligning them using DALI against SIX effector representatives from each family to assess if they fell into any of the established families (Fig 3B). We found the predicted structure of FOXGR_015533 adopts a nine β-stranded sandwich and is likely a member of the ToxA-like class (Fig 3C). PSL1 [17] and FOXGR_015322, here designated PSL2, are sequence related effectors (∼85% sequence identity) and show a conserved structure (Fig 3E). Both have Z-scores of >2 against Family 4 and are likely members of this family.

Based on this analysis we also suggest an additional structural family. FOXG_18699 and FOXGR_015522 are structurally related (Z-score of 2.2) with a sequence identity of ∼29%. While FOXGR_015522 does share some resemblance to Family 4, based on manual alignment (Fig 3F) and domain topology analysis (S8 Fig) these effectors appear to belong to an independent structural family, designated Family 5. Collectively, these data demonstrate that *Fol* utilises multiple structurally related, sequence diverse, effectors during infection of tomato.

### Interaction between effector pairs from two structural families

In *Fol, Avr2* and *SIX5*, and *SIX8* and *PSL1* form a similar head-to-head relationship in the genome with shared promoters and are divergently-transcribed (Fig 4A) [16, 17]. Previously, studies concerning Avr2 and SIX5 have demonstrated that the proteins function together and interact directly via yeast-two-hybrid analysis [9]. Homologues of SIX8 and PSL1 from *Focn* (SIX8 and PSE1) are also functionally dependent on each other, however an interaction could not be established in yeast [17]. Here we demonstrate that both protein pairs contain a ToxA-like family member (Avr2, SIX8) and a structural family 4 member (SIX5, PSL1). Considering the predicted structural similarities, we were interested in testing whether *Fol* SIX8 and PSL1 interact.

**Fig 4.**
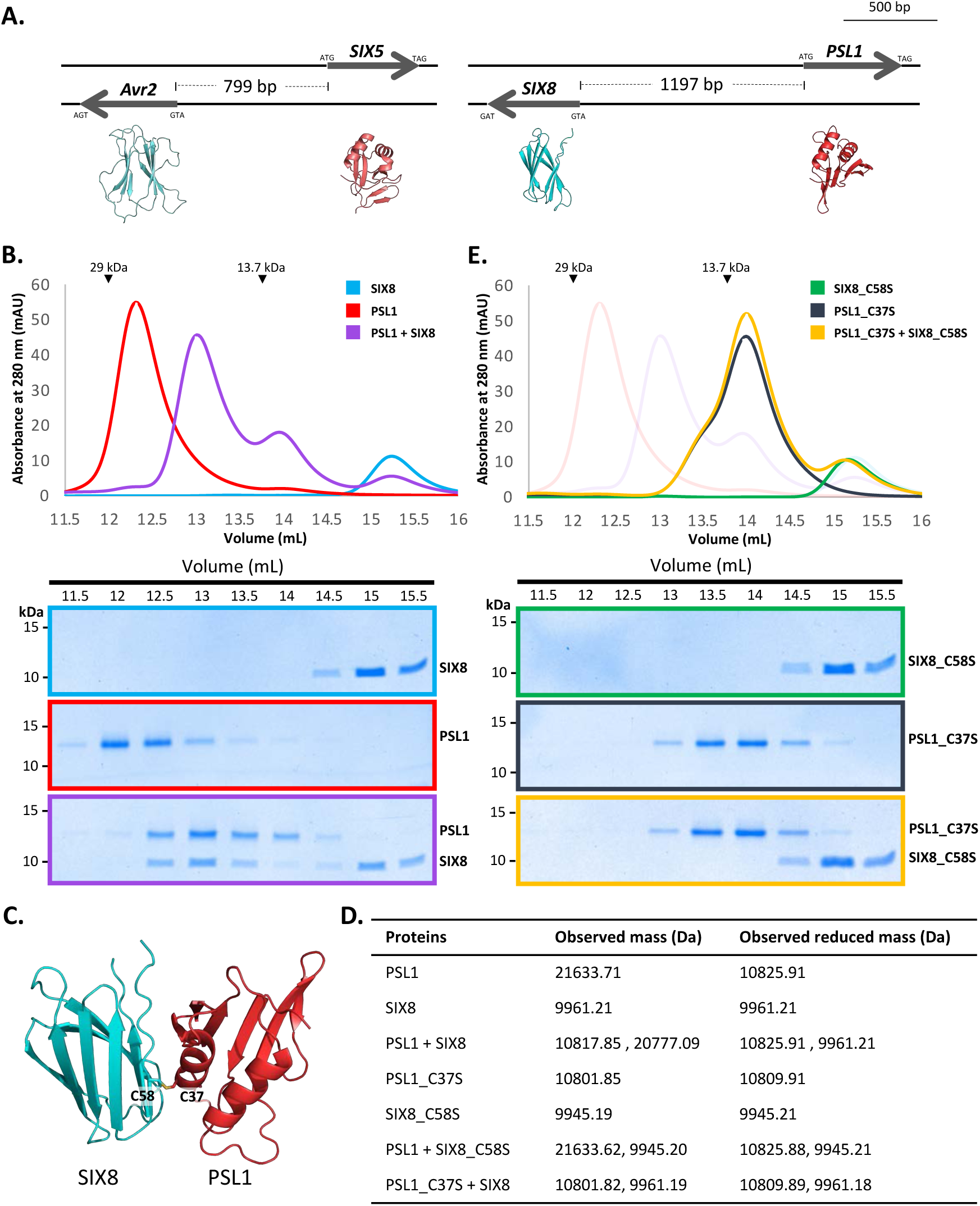
PSL1 and SIX8 interact *in vitro* mediated by an intermolecular disulfide bond. **(A)** Schematic representation of the *Avr2 (SIX3)* – *SIX5* and *SIX8* – *PSL1* loci within *Fol*. AlphaFold2 models or experimentally solved protein structures are shown underneath. **(B)** Size exclusion chromatograms of PSL1 alone (red), SIX8 alone (blue), PSL1 and SIX8 (purple) (following a 30 min incubation) separated over a Superdex S75 Increase SEC column (top panel). Equal concentrations of the protein were used (note the absorbance of SIX8 at 280 nm is ∼0.3 resulting in a smaller absorbance and peak height). Indicated sizes above the chromatogram are based on protein standards run under similar conditions as presented in the manufacturer’s column guidelines. Coomassie-stained SDS-PAGE gels depicting samples taken from 500 µL fractions corresponding to the volumes indicated above the gels, with molecular weights (left) and proteins (right) annotated (bottom panels). **(C)** Model of the SIX8-PSL1 complex generated by AlphaFold2-Multimer (top model shown). Co-localisation of Cys 58 from SIX8 and Cys 37 from PSL1 shown in stick form. **(D)** Observed masses of PSL1 and SIX8 protein mixtures by intact mass spectrometry (MS). Samples were treated with or without the reducing agent DTT prior to MS. The deconvoluted mass spectra of all proteins can be found in S9 Fig. **(E)** As for **(B)** but with PSL1_C37S (black), SIX8_C58S (green), and PSL1_C37S and SIX8_C58S (yellow).

We heterologously produced *Fol* SIX8^50-141^ and PSL1^18-111^ (S1E-F Fig) (after referred to as SIX8 and PSL1) and co-incubated the proteins before analysing by size exclusion chromatography (SEC) (Fig 4B). The elution profile of PSL1 alone showed a major peak (∼12.25 mL) at a volume consistent with a dimeric form of the protein, while SIX8 showed a major peak (∼15 mL) consistent with a monomer (Fig 4B). Strikingly, when incubated together the major protein peaks migrate to ∼12.8 mL. SDS-PAGE analysis confirmed that presence of PSL1 and SIX8, indicating that the migration of both proteins on SEC is altered after incubation (Fig 4B). These data are consistent with PSL1 and SIX8 forming a heterodimer.

To understand the structural basis of the interaction, we attempted to solve the structure of the complex, but we were unable to obtain crystals. We subsequently utilised AlphaFold2-Multimer [46] through ColabFold [47], to model the interaction. Manual inspection of the top 5 models (S10A Fig, top model shown Fig. 4C) demonstrated that the thiol side chain of a free cysteine in PSL1 (Cys 37) and SIX8 (Cys 58) co-localised in the dimer interface, suggesting that an inter-disulfide bond may mediate the interaction. To test this, we performed intact mass spectrometry of SIX8 and PSL1 (alone and post incubation) under non-reduced and reducing conditions. The mass observed from the incubated SIX8 and PSL1 non-reduced sample contained a predominant species consistent with the combined molecular weight of SIX8 and PSL1 (20777 Da) (Fig 4D, S9G-H Fig). SIX8 and PSL1 failed to form a heterodimer with an unrelated protein containing a free cysteine, suggesting specificity in the interaction (S9I-L Fig). Collectively, these data demonstrated that the SIX8-PSL1 heterodimer is mediated via a disulfide bond.

To confirm the involvement of the predicted residues involved, interaction with cysteine mutants of PSL1 and SIX8 (PSL1_C37S^18-111^ and SIX8_C58S^50-141^, after referred to as PSL1_C37S and SIX8_C37S) were analysed (Fig 4E). When PSL1_C37S was incubated with SIX8_C37S or SIX8 alone, the heterodimer was not resolved via SEC (Fig 4D, S10B Fig). This was further confirmed using mass spectrometry (Fig 4C). We crystallised and solved the structure of SIX8_C58S at 1.28 Å (S1E Fig and S10C Fig) which confirms its inclusion within the ToxA-like structural family (S10D Fig).

### The molecular basis of Avr1 recognition by the I receptor

The structural identification of the FOLD effector family provides an opportunity to understand their recognition by cognate immunity receptors. Here, we focussed on Avr1 (SIX4), which is recognised by the I immunity receptor leading to ETI and disease resistance [19]. Previous studies have shown co-expression of the *I* gene from the M82 tomato cultivar (I^M82^) with Avr1 in *Nicotiana benthamiana* leads to a cell death response, a proxy for ETI [19]. Conversely, co-expression with the allelic variant (i^Moneymaker^) from the susceptible cultivar Moneymaker does not lead to cell death as the receptor cannot recognise Avr1 [19] (Fig 5B). Here we sought to further define the recognition between Avr1 and I utilising the *N. benthamiana* system.

**Fig 5.**
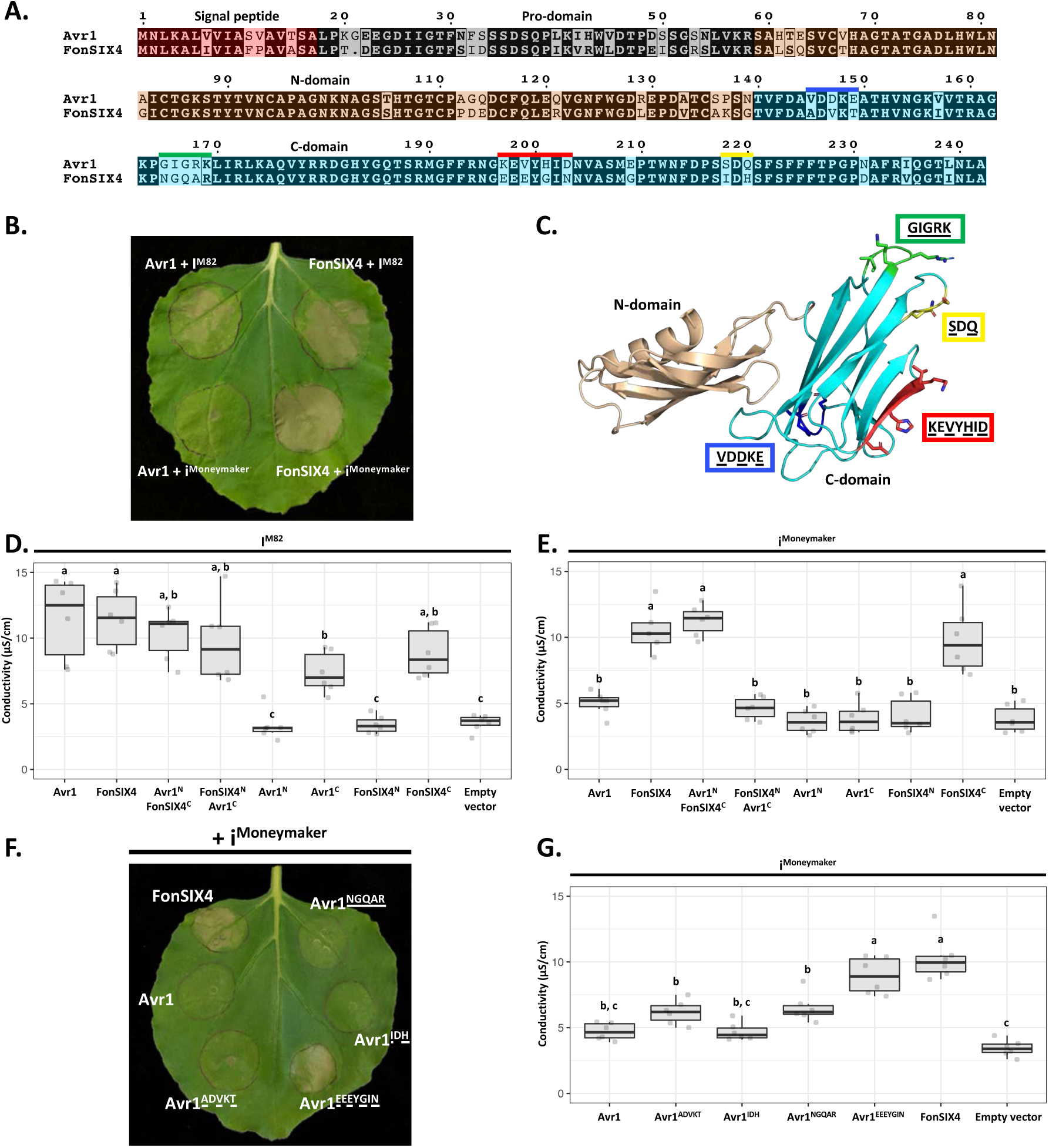
The C-domain of Avr1 mediates recognition by the I receptor. **(A)** Amino acid sequence alignment of Avr1 and FonSIX4, a homologue from *F. oxysporum* f. sp. *niveum*. The signal peptide, pro-domain, N-domain and C-domain are highlighted in red, grey, beige and blue, respectively. Within the C-domain, surface exposed regions that differ between Avr1 and FonSIX4 are overlined. **(B)** Avr1 and FonSIX4 were transiently expressed in *Nicotiana benthamiana* with either I^M82^ or i^Moneymaker^ via *Agrobacterium-*mediated transformation (n = 6). **(C)** The crystal structure of Avr1, showing the N-and C-domains in beige and light blue, respectively as represented in A. Regions containing variant residues within the C-domain between Avr1 and FonSIX4 are coloured corresponding the overlined colours in A. Variant residues are underlined and represented in stick form. **(D)** Ion leakage conductivity of the Avr1 and FonSIX4 chimeric constructs, and N-and C-domains individually, when transiently co-expressed with I^M82^ or **(E)** i^Moneymaker^. Two additional independent experiments were repeated with similar results (S12 Fig). **(F)** Leaf image and **(G)** ion leakage quantification of the Avr1 mutants (Avr1<u>^A^</u>^D<u>V</u>KT^, Avr1<u>^NGQAR^</u>, Avr1^ID<u>H</u>^, Avr1^EEEY<u>GIN</u>^) when transiently co-expressed with i^Moneymaker^ (n = 6). Variant residues between Avr1 and FonSIX4 are underlined. Six biological replicates for each construct were measured using an ion leakage assay. One-way ANOVA and post *hoc* Tukey’s honestly significant difference tests were performed. Treatments that do not share a letter are significantly different from each other at *p < 0.05.* Leaves were imaged 5 days post infiltration (dpi).

To facilitate this, we identified homologues of Avr1 that possess natural residue variation. FonSIX4, a homologue of Avr1 from the watermelon pathogen, *F. oxysporum* f. sp. *niveum* (*Fon*) shares 79% identity with Avr1 (Fig 5A). Using the *N. benthamiana* assay we show FonSIX4 is recognised by I receptors from both cultivars (I^M82^ and i^Moneymaker^) (Fig 5B). FonSIX4 and Avr1 differ by 34 residues distributed across both N-and C-domains of the protein (Fig 5A). To narrow down the regions involved in recognition we designed chimeric variants by swapping the N-and C-domains (Avr1^N^FonSIX4^C^ and FonSIX4^N^Avr1^C^) (Fig 5A, C). When these were co-expressed with i^Moneymaker^ the cell death response, quantified using ion leakage assays (Fig 5D-E) and visual inspection (S11A Fig), suggest the C-domain of FonSIX4 is recognised by i^Moneymaker^. We separated Avr1 and FonSIX4 proteins into their N-or C-domains and co-expressed with I^M82^ or i^Moneymaker^. Quantification using ion leakage assays demonstrate that the C-domains of Avr1 and FonSIX4 cause cell death when expressed with I^M82^ and I^M82/Moneymaker^, respectively. These data confirm the C-domain is sufficient for I receptor recognition (Fig 5D-E, S12 Fig, see S11 Fig for *N. benthamiana* leaf infiltration and protein accumulation data).

To understand how Avr1 can escape i^Moneymaker^ recognition, we focussed on surface exposed variant residues (underlined) mapping to four regions within the C-domain (Fig 5A and 5C). Four reciprocal swap mutants between Avr1 and FonSIX4 (Avr1^AD<u>V</u>KT^, Avr1^ID<u>H</u>^, Avr1<u>^NGQAR^</u>, Avr1^EEEY<u>GIN</u>^) were co-expressed with i^Moneymaker^ to identify the residues required for FonSIX4 recognition. Avr1^EEEY<u>GIN</u>^ showed consistent ion leakage and cell death similar to FonSIX4 (Fig 5F-G), whereas ion leakage quantification for the other three mutants (Avr1^AD<u>V</u>KT^, Avr1^ID<u>H</u>^, Avr1<u>^NGQAR^</u>) was statistically similar to the non-recognised Avr1 (Fig 5G). The reciprocal mutations in FonSIX4 (FonSIX4^KE<u>V</u>Y<u>HID</u>^) significantly reduced ion leakage and cell death response when co-expressed with i^Moneymaker^ compared to FonSIX4 (S11D-E Fig, see S11G Fig for protein accumulation data). Collectively, these data show that the C-domain in Avr1 is recognised by I^M82^, and surface exposed residues in the C-domain allow Avr1 to escape recognition by i^Moneymaker^.

## Discussion

Pathogenic fungi are in a continuous arms race with their plant hosts. To aid virulence, but avoid detection, effectors evolve rapidly causing significant diversity at the amino acid sequence level [48]. An emerging theme in fungal effector biology is the classification of effectors into families based on structural similarity [49]. Here, we demonstrate that despite their sequence diversity, the *Fol* SIX effectors can be classified into a reduced set of structural families. This observation has implications for functional studies of SIX effectors, and ultimately our understanding of the infection strategies used by *F. oxysporum*.

### Expanding the structural classes in fungal effectors

To date, five fungal effector families have been defined based on experimentally-determined structural homology, including the MAX [27], RALPH [28, 50, 51], ToxA-like [24-26], LARS [29, 52] and FOLD effectors, defined here. Effectors that fall within many of these structural families are shared across distantly related fungal species. The ToxA-like family includes effectors from fungi that group to both divisions of higher-fungi (Basidiomycota and Ascomycota fungi) [24-26]. The MAX effector family were originally defined as AVR effectors from *M. oryzae* and ToxB from *P. tritici-repentis* [27] but pattern-based sequence searches suggest they are widely distributed amongst the Dothideomycetes and Sordariomycetes [27, 53]. Similarly, LARS effectors, defined in *Leptosphaeria maculans* and *Fulvia fulva,* have structural homologues predicted in at least 13 different fungal species [29]. Based on sequence homologues alone, FOLD effectors are well dispersed in fungi with homologues amongst the Sordariomycetes including many *formae speciales* of *F. oxysporum*, *Colletotrichum* and *Ustilaginoidea*. Based on structural comparison of the AlphaFold2 structural database we show that is extended to fungi in three Divisions, including plant pathogens and symbionts. This was supported by a recent study modelling the secretomes of arbuscular mycorrhizal fungi which found enlarged and diversified gene families encoded proteins predicted to share the FOLD effector structure [54]. The exclusive presence of FOLD effectors in plant-colonising fungi may suggest they facilitate plant colonisation in pathogenic and symbiotic fungi [54].

### Effector structure prediction

Experimentally determining the structures of fungal effectors is not a trivial undertaking. From challenges associated with effector protein production through to hurdles related to structure solution (such as experimental phasing), the research time required to determine an effector structure experimentally ranges from months to many years (sometimes never). Not surprisingly, any reliable structural modelling methods are welcomed by researchers interested in effector biology. To this end, several recent studies have used effector structure prediction to expand our understanding of plant-microbe interactions [55, 56].

Work by Bauer and colleagues, prior to the release of AlphaFold2, used structural modelling to show that numerous recognised Avr effectors from the barley powdery mildew-causing fungal pathogen *Blumeria graminis* (*Bgh*) are members of the RALPH effectors class [55]. Seong and Krasileva used similar structural modelling approaches to predict the folds of ∼70% of the *Magnaporthe oryzae* secretome [56]. In doing so, they suggested an expansion in the number of MAX effectors and identified numerous sequence-unrelated groups of structural homologues (putative structural classes) within *M. oryzae*. Making use of AlphaFold2, Yan and colleagues show that structurally conserved effectors, including the MAX effector family, from *M. oryzae* are temporally co-expressed during the infection process [57]. In the largest comparison study to date, Seong and Krasileva carried out a large comparative structural genomics study of fungal effectors utilising AlphaFold2 [58]. Their findings support the hypothesis that the structurally conserved effector families are the result of divergent evolution and support previous finding that the structural landscape of effectors is more limited than what is suggested by sequence-diversification.

Here, we were in a unique position to apply and benchmark AlphaFold2 against experimentally determined structures for *Fol* effector prediction. We subsequently used AlphaFold2 to demonstrate that, within the repertoire of effectors we tested, up to five sequence-unrelated structural families are secreted during *Fol* infection. There are numerous caveats in relying solely on AlphaFold2 to generate structural models of effectors. The accuracy of models generated by AlphaFold2 can decline in cases with low numbers of homologues (∼30 sequences in the multiple sequence alignment) [34]. This may help explain the low confidence prediction for SIX4 (Avr1) (S4A Fig), which is only distributed in a few *ff. spp.* of *F. oxysporum*. This poses a potential issue for predicting the structures of fungal effectors that lack homologues. In our hands, we have had mixed results when comparing several unpublished effector structures experimentally determined in our lab to AlphaFold2 models. In some instances, the models are wrong, for example AvrSr50 [59], however, in these cases the AlphaFold2 predictions reported low confidence scores, an important criterion for assessment of model reliability. Despite this, AlphaFold2 models were critical in solving the structure of SIX6 and SIX8, as templates for molecular replacement. This negated the need to derivatise our crystals, a process that we had struggled with for SIX6 crystals, significantly reducing the time and research effort to determine the experimental structures.

### Structural classes: A starting point for functional characterisation

Given their lack of sequence identity to proteins of known function or conserved motifs, structural determination of effectors is often pursued to provide functional insight and understanding of residues involved in recognition. The existence of structural families of effectors raises the question of whether links can now be made concerning their function based on structural similarities. Unfortunately, the FOLD effectors share little overall structural similarity with known structures in the PDB outside of the similarity with each other. At a domain level, the N-domain of the FOLD effector Avr1 has some structural similarities with cystatin cysteine protease inhibitors (PDB code: 4N6V, PDB code: 5ZC1) [60, 61], and the C-domain with tumour necrosis factors (PDB code: 6X83) [62] and carbohydrate-binding lectins (PDB code: 2WQ4) [63]. Relatively weak hits were observed for Avr3/Six6.

Interestingly, the predicted models for SIX9 and SIX11 within Family 3 have structural homology with RNA-binding proteins (PDB code: 3NS6, PDB code: 5X3Z) [42, 64], unrelated to RALPH effectors. Despite this structural homology, close inspection of these models suggests RNA binding is unlikely, as in both models the putative RNA binding surface is disrupted by a disulfide bond.

The putative family 4 effectors (SIX5, SIX14, PSL1 and PSL2) have structural homology with KP6 effectors and heavy metal associated (HMA) domains. Metal binding within HMA domains is facilitated by conserved cysteine residues [65], however, their absence in the family 4 effectors suggests they are unlikely to have this activity.

The putative family 5 effectors (FOXGR_015522 and FOXG_18699) have structural homology with different proteins within the PDB. FOXGR_015522 is structurally similar to plant defensins (PDB code: 6MRY, PDB code: 7JN6) [66, 67] and K^+^ channel-blocking scorpion toxins (PDB code: 1J5J, PDB code: 2AXK) [68, 69]. FOXG_18699 has structural homology with the C-terminal domain of bacterial arginine repressors (PDB code: 1XXB, PDB code: 3CAG) [70, 71].

### A structural explanation for functional effector pairs

One interesting outcome of this study is a link between structural families and co-operative interactions between effectors. The ToxA-like effectors, Avr2 and SIX8 are known to form functional effector pairs with SIX5 and PSE1 (PSL1-homolouge), respectively [9, 17]. According to our modelling work, both SIX5 and PSL1 are members of structural family 4. *Avr2* and *SIX5* are adjacent divergently-transcribed genes on *Fol* chromosome 14 and the protein products have been shown to physically interact [9]. Likewise, *SIX8* and *PSL1* are adjacent divergently-transcribed genes in the *Fol* genome and we demonstrate here a physical interaction between the proteins. The AlphaFold2-multimer models of the SIX8 and PSL1 heterodimer, drew our attention to the inter-disulfide bond between SIX8 and PSL1 required for the interaction, which we confirmed experimentally. While these residues are conserved in *Focn* SIX8 and PSE1, the Avr2 structure and SIX5 model lack free cysteine residues, suggesting a different mode of interaction.

Interestingly, two other *SIX* genes also form a divergently-transcribed gene pair on *Fol* chromosome 14. *SIX7* (ToxA-like family) and *SIX12* possess start codons 2,319 base-pairs apart and potentially share a common promoter. While SIX12 did not group with any structural families, the AlphaFold2 model had a very low prediction confidence (35.5%). On closer inspection of the sequence, we observed that the cysteine spacing in SIX12 closely resembles other family 4 members (S13 Fig), which suggests that SIX12 may also be a family 4 member. We therefore speculate that SIX7 and SIX12 may function together, as described for the Avr2/SIX5 and SIX8/PSL1 pairs.

### Are experimentally derived effector structures still worth the effort?

The potential of machine-learning structural-prediction programs, such as AlphaFold2, heralds an exciting era, especially for a field that has long suffered from a lack of prediction power based on effector sequences. A question now emerges; when prediction model confidence is high, should we bother solving structures experimentally? The answer to such a question will always depend on what the structure is being used for. Ultimately, structural models, whether experimentally or computationally derived, represent information to base and/or develop a hypothesis to subsequently test. Here we demonstrate the power of structure prediction in combination with experimentation, both for validating models and understanding protein:protein interaction interfaces. One interesting observation we made was that while the AphaFold2-multimer models of the SIX8 and PSL1 heterodimer were sufficient to highlight the cysteine residues required for mediating the interaction, the models and interaction interfaces differed significantly (S10A Fig). When the modelling was repeated with the SIX8^C58S^ experimentally derived structure included as a template, the interaction models and heterodimer interface were of higher quality and essentially identical (S10E Fig). This observation can be retrospectively reconciled. The region of SIX8 involved in the interaction with PSL1 was modelled incorrectly by AlphaFold2 when compared to the structure (S10D Fig). Collectively, these data highlight that some models are good enough, but others maybe better.

### Effector structural classes and understanding receptor recognition

Understanding the structural basis of plant immunity receptor-effector interactions represent a key step towards engineering plant immunity receptors with novel specificities. Recent structures of nucleotide-binding domain leucine-rich repeat (NLR) proteins reveal exquisite details of these direct interactions [72-74]. The FOLD effectors, Avr1 and Avr3, are recognised by different classes of immunity receptors; I, an LRR-RP [19] and I-3, a SRLK [21]. While the mode of recognition has not yet been described for Avr3, we demonstrate here that Avr1 is recognised at the C-domain (Fig 5). This is significant because it demonstrates that different immunity receptor classes can recognise structural homologues. It might also help explain the function of Avr1 during *Fol* infection. When Houterman and colleagues identified Avr1, they demonstrated that it could suppress plant immunity conferred by the *I-2* and *I-3* receptors [22]. Considering our structural understanding of these FOLD effectors, it is plausible that Avr1 achieves suppression of I-3-mediated immunity by preventing Avr3 recognition through competitive inhibition. The LARS effectors represent another example of effectors that can activate and suppress resistance-gene-mediated immunity. AvrLm4-7 can prevent recognition of AvrLm3 and AvrLm9 (all LARS structural homologues [29]), by their cognate Rlm receptors [75, 76]. *Rlm4*, *Rlm7* and *Rlm9* all encode for wall-associated kinases [77, 78], but the identify of *Rlm3* remain unknown. These studies demonstrate that members of at least two different structural effector families can suppress immunity triggered by structurally homologous effectors.

Collectively, the results presented here will aid future studies to understand the molecular basis of *F. oxysporum* effector function and recognition, and by extension, the design and engineering of immunity receptors with novel recognition specificities to help protect plants against *Fusarium* wilt disease.

## Materials and methods

### Vectors and gene constructs

SIX6, Avr1Thrombin, SIX6-TEV, SIX8Thrombin, SIX8_C58SThrombin, PSL1, PSL1_C37S and SIX13 coding sequences (without their signal peptides as determined by SignalP-5.0) were codon optimised for expression in *E. coli* and synthesised with Golden-Gate compatible overhangs by Integrated DNA Technologies (IDT, Coralville, USA) (S4 Table). The Kex2 cleavage motif of Avr1 and SIX8 were replaced with a thrombin cleavage motif, and TEV protease cleavage motif for SIX6 for pro-domain processing. Avr1 and Avr3 coding sequences were PCR amplified using *Fol* cDNA as a template with primers containing Golden-Gate compatible overhangs. All genes for *E. coli* expression were cloned into a modified, Golden-Gate-compatible, pOPIN expression vector [79]. The final expression constructs contained N-terminal 6xHis-GB1-tags followed by 3C protease recognition sites. The Golden-Gate digestion, ligation reactions and PCR were carried out as described by Iverson, Haddock [80]. Avr1 and FonSIX4 mutant sequences without the signal peptide were synthesised with compatible overhangs by IDT (S4 Table) and cloned into the pSL vector containing the *Nicotiana tabacum* PR1 signal peptide using the In-fusion cloning kit (Takara Bio USA Inc., San Jose, USA) to allow efficient secretion in *N. benthamiana* via *Agrobacterium*-mediated expression. For tagged constructs, Avr1 and FonSIX4 mutant sequences and 3xHA tag were amplified with PCR and assembled using In-fusion cloning into the pSL vector. All of the primers were synthesised by IDT (S5 Table). All constructs were verified by sequencing.

### Protein expression and purification

Sequence-verified constructs were co-expressed with CyDisCo in SHuffle T7 Express C3029 (New England Biolabs (NEB), Ipswich, USA) and purified as previously described [32]. For Avr3, the buffers used after fusion tag cleavage were altered slightly to increase protein stability and a second IMAC step was excluded after the cleavage of the N-terminal fusion tag. During the cleavage step, the protein was dialysed into a buffer containing 10 mM MES pH 5.5 and 300 mM NaCl. The size-exclusion chromatography (SEC) HiLoad 16/600 Superdex 75 pg column (Cytiva) was equilibrated with a buffer containing 10 mM MES pH 5.5 and 150 mM NaCl.

For biochemical and crystallisation studies, Avr1 and SIX8 with an internal thrombin cleavage site, and SIX6 with an internal TEV protease cleavage site for pro-domain removal were processed with 2 to 4 units of thrombin from bovine plasma (600-2,000 NIH units/mg protein) (Sigma-Aldrich Inc., St. Louis, USA) per mg of protein at 4°C or TEV protease (produced in-house) until fully cleaved. Fully cleaved proteins were purified further by SEC using a HiLoad 16/600 or HiLoad 26/600 Superdex 75 pg column (Cytiva) equilibrated with a buffer containing 10 mM HEPES pH 7.5 or 8.0 and 150 mM NaCl. Proteins were concentrated using a 10 or 3 kDa molecular weight cut-off Amicon centrifugal concentrator (MilliporeSigma, Burlington, USA), snap-frozen in liquid nitrogen and stored at -80°C for future use.

### Intact mass spectrometry

Proteins were adjusted to a final concentration of 6 µM in 0.1% (v/v) formic acid (FA) for HPLC-MS analysis for untreated samples. For reduced samples, DTT was added to the protein to a final concentration of 10 mM. Proteins were incubated at 60°C for 30 minutes and adjusted to 6 µM in 0.1% (v/v) FA. Intact mass spectrometry on all proteins was carried out as described previously [32]. Data were analysed using the Free Style v.1.4 (Thermo Fisher Scientific) protein reconstruct tool across a mass range of m/z 500 – 2000 and compared against the theoretical (sequence based) monoisotopic mass.

### Circular dichroism (CD) spectroscopy

The CD spectra of purified effectors of interest were recorded on a Chirascan spectrometer (Applied Photophysics Ltd., UK) at 20°C. Samples were diluted to 10 µM in a 20 mM sodium phosphate buffer at pH 8.0. Measurements were taken at 1 nm wavelength increments from 190 nm to 260 nm. A cell with a pathlength of 1 mm, a bandwidth of 0.5 nm and response time of 4 s were used, with 3 accumulations. The data were averaged and corrected for buffer baseline contribution, and visualised using the webserver CAPITO tool with data smoothing [81]. CD analysis was performed on all purified proteins (S14 Fig).

### Crystallisation, diffraction data collection and crystal structure determination

Initial crystallisation screening was performed for Avr3^22-284^, Avr1^18-242^, Avr1^59-242^, SIX8^50-141^, PSL1^18-111^, SIX6^17-225^, SIX6^58-225^, SIX8_C58S^19-141^, SIX8_C58S^50-141^, PSL1_C37S^18-111^, SIX8-PSL1 complex and SIX13^22-293^ with and without Kex2 protease using 150 nL protein solution and 150 nL reservoir solution sitting-drop plates at 18°C with commercially available sparse matrix screens.

No crystals were obtained for Avr1^18-242^, SIX6^17-225^, SIX8^50-141^, PSL1^18-111^, SIX8-PSL1 complex and SIX13^22-293^. Final crystallisation conditions were optimised for Avr3^22-284^ (0.2 M ammonium sulfate, 0.1 M Bis-Tris pH 6.5, 25% (w/v) PEG 3350), Avr1^59-242^ (0.2 M ammonium sulfate, 0.1 M sodium acetate pH 4.5, 17.5% (w/v) PEG 4000), SIX6^58-225^ (0.2 M ammonium tartrate and 20% (w/v) PEG 3350), SIX8_C58S^50-141^ (0.17 M ammonium sulfate, 15% (v/v) glycerol and 25.5% (w/v) PEG 4000), SIX13 (0.2 M lithium sulfate, 0.1 M Bis-Tris pH 6.5, 25% (w/v) PEG 3350) and PSL1_C37S^18-111^ (70% (v/v) MPD and 0.1 M HEPES pH 7.5). Detailed crystallisation optimisation can be found in the supplementary methods.

Crystals were transferred into a cryoprotectant solution containing reservoir solution and 15% (v/v) ethylene glycol, 20% (v/v) glycerol or 10% (v/v) ethylene glycol and 10% (v/v) glycerol. No cryoprotecting was required for SIX8_C58S^50-141^ and PSL1_C37S^18-111^ crystals as the conditions contained sufficient cryoprotectant (glycerol and MPD, respectively) within the crystallisation condition. For experimental phasing, Avr3^22-284^ and Avr1^59-242^ crystals were soaked in a cryoprotectant solution containing 0.5 M or 1 M sodium bromide and vitrified in liquid nitrogen. The datasets for bromide-soaked crystals were collected on the MX1 beamline at the Australian Synchrotron [82] (S1 Table). The datasets were processed in XDS [83] and scaled with Aimless in the CCP4 suite [84, 85]. The CRANK2 pipeline in CCP4 was used for bromide-based SAD phasing [86, 87]. Models were refined using phenix.refine in the PHENIX package [88] and model building between refinement rounds was done in COOT [89]. The models were used as a template for molecular replacement against high resolution native datasets collected on the MX2 beamline at the Australian Synchrotron [90]. Automatic model building was done using AutoBuild [91], and subsequent models were refined with phenix.refine and COOT. For SIX6^58-225^ and SIX8_C58S^50-141^, high confidence *ab initio* models were generated with AlphaFold2 (S3 Fig), which was used as a template for molecular replacement against a native dataset collected on the MX2 beamline at the Australian Synchrotron. The resultant structure was refined as described above.

### Structural modelling and structural alignment

Structural models were generated with Google DeepMind’s AlphaFold2 using the amino acid sequences of SIX effectors and candidates without the signal peptide, as predicted by SignalP-5.0 [92] and predicted pro-domain by searching for a Kex2 cleavage motif (KR, RR or LxxR) if present [30] (S3 Table; S6 Fig). For AlphaFold2 predictions the full databases were used for multiple sequence alignment (MSA) construction. All templates downloaded on July 20, 2021 were allowed for structural modelling. For each of the proteins, we produced five models and selected the best model (ranked_0.pdb). Pairwise alignments of the structural models generated by AlphaFold2 and the experimentally determined structures of Avr1 (PDB code: 7T6A), Avr3 (PDB code: 7T69), SIX6 (PDB code: 8EBB) and SIX8 (PDB code: 8EB9) were generated using the DALI server all against all function [33]. Structural similarity between the pairwise alignments were measured using Z-scores from the DALI server.

### Distribution of FOLD family members across fungi

Structure based searches to determine the distribution of FOLD effectors across other phytopathogens was carried out by searching the experimentally determined Avr1, Avr3 and SIX6 structures against available structure databases (Uniprot50, Proteome, Swiss-Prot) using the Foldseek webserver [41] using a 3Di search limited to fungi. An e-value cut off of 0.01 was used, and non-plant associated fungi were removed as well as duplicated results for final analysis. Proteins below 100 amino acids, and above 500 amino acids were filtered out and remaining structural hits were manually inspected for similarity to FOLD effectors.

### Interaction studies between PSL1 and SIX8

To investigate the PSL1 and SIX8 interaction *in vitro*, ∼140 µg of PSL1^18-111^ and SIX8^50-141^ individually, and ∼140 µg PSL1^18-111^ and 140 µg of SIX8^50-141^ together were injected onto a Superdex 75 Increase 10/300 (Cytiva) column pre-equilibrated in 20 mM HEPES pH 7.5, 150 mM NaCl, after a 30 min room temperature incubation. To investigate the residues responsible for the interaction, SIX8_C58S^50-141^ and PSL1_C37S^18-111^ mutants were used instead. Samples across the peaks were then analysed by Coomassie-stained SDS-PAGE. To investigate the mode of interaction, PSL1 and SIX8 proteins and mutants at 10 µM were incubated individually or together for 1 hour at room temperature. An unrelated protein with a free cysteine (AvrSr50^RKQQC^) [59] was used to assess the specificity of the PSL1-SIX8 interaction. Proteins were analysed by intact mass spectrometry with or without the addition of DTT as described above.

### *Agrobacterium*-mediated gene expression in *N. benthamiana*

*Agrobacterium tumefaciens* cultures containing the pSL constructs and pSOUP [93] were diluted to an OD600 of 1.0 in 10 mM MES pH 5.5 buffer containing 10 mM MgCl2 and 200 μM acetosyringone and incubated in the dark for 2 hours. For co-infiltrations, cultures were mixed together in equal volumes. Resuspensions were infiltrated into 4 – 5-week-old *N. benthamiana* leaves. Infiltrated plants were kept in a 25°C controlled temperature room with a 16-hour photoperiod. Leaves were imaged 4 – 7 dpi.

### Ion leakage assay

Six biological replicates each consisting of three leaf discs (7 mm diameter) were harvested from leaves infiltrated with *Agrobacterium* 20 - 24 hours post infiltration and incubated in 7 mL of water in a 6-well culture plate. The water was replaced after 40 - 60 min. The leaf discs were incubated in water at room temperature and the conductivity was measured after 24 - 48 hours.

### Immunoblot analysis of proteins expressed in *N. benthamiana*

*N. benthamiana* leaves infiltrated with *A. tumefaciens* cultures were harvested 3 dpi. Leaf tissue was frozen in liquid nitrogen, ground into a powder and resuspended in 3x Laemmli buffer containing 0.2 mM DTT and 5 M urea to extract proteins. Samples were boiled for 10 min and centrifuged at 13000 xg to remove leaf debris. Proteins were separated by SDS-PAGE and transferred by electroblotting onto PVDF membranes. Protein blots were probed with anti-HA antibodies conjugated to horseradish peroxidase (Roche, Switzerland, 12013819001, 1:4000). Immunoblots were visualised with Pierce ECL Plus Western Blotting Substrate (Thermo Fisher Scientific) as described by the manufacturer. Membranes were stained with Ponceau S to assess protein loading.

## Acknowledgements

This work was supported by the Australian Research Council (ARC DP200100388 D.J./S.W.) and the Australian Academy of Science (Thomas Davies Grant). S.W. was funded by an ARC Future Fellowship (FT200100135) and supported by the ANU Future Scheme (35665). L.M. was funded by an ARC Discovery Early Career Researcher Award (DE170101165). A.S. was a recipient of the AINSE Honours Scholarship Program, and D.Y. and C.M. held an AINSE Postgraduate Research Award. P.K. was supported by a Netaji Subhas ICAR International Fellowship. The authors acknowledge the use of the ANU crystallisation facility. This research was undertaken in part using the MX2 beamline at the Australian Synchrotron, part of ANSTO, and made use of the Australian Cancer Research Foundation (ACRF) detector. The authors acknowledge use of the Australian Synchrotron MX facility and thank the staff for their support. The coordinates and structure factors for Avr1, Avr3, SIX6 and SIX8 have been deposited in the Protein Data Bank with accession number 7T6A, 7T69, 8EBB and 8EB9, respectively.

## Supplementary Figures

**S1 Fig.**
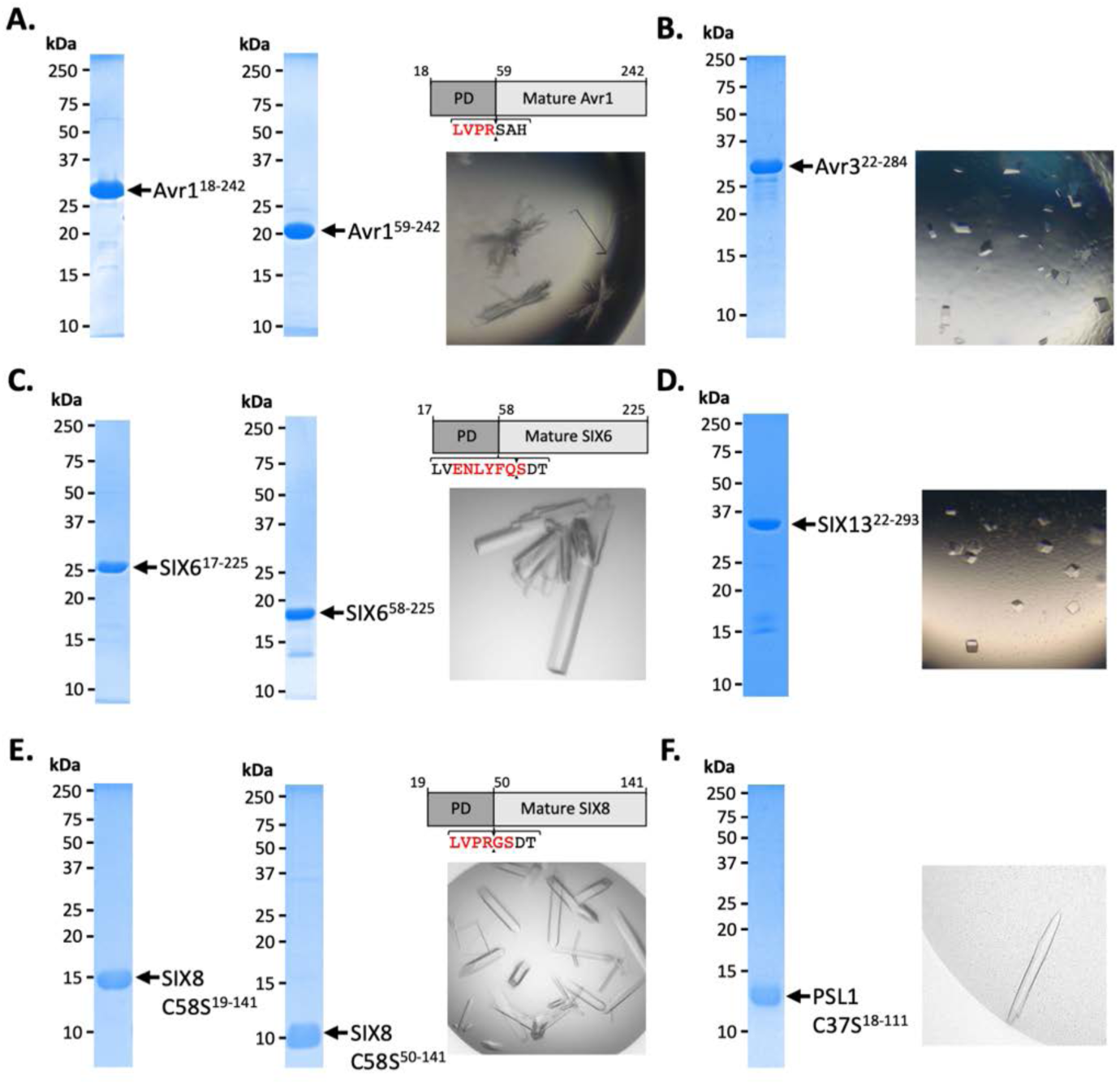
Crystallisation of Avr1, Avr3, SIX6, SIX8, SIX13 and PSL1. **(A)** Coomassie-stained gel showing Avr1^18-242^ (left panel) and mature Avr1^59-242^ cleaved *in vitro* with thrombin (middle panel). Schematic of Avr1 engineered with an internal thrombin cleavage site replacing the Kex2 cleavage motif after the pro-domain (PD) (top right panel). Optimised crystals of Avr1^59-242^ (bottom right panel) **(B)** Coomassie-stained gel showing purified Avr3^22-284^ used for crystallisation studies (left panel). Optimised crystals of Avr3 (right panel). **(C)** Coomassie-stained gel showing SIX6^17-225^ (left panel) and mature SIX6^58-225^ cleaved *in vitro* with TEV protease (middle panel). Schematic of SIX6 engineered with an internal TEV protease cleavage site replacing the Kex2 cleavage motif after the PD (top right panel). Optimised crystals of SIX6^58-225^ (bottom right panel). **(D)** Coomassie-stained gel showing SIX13^22-293^ protein (left panel). Optimised crystals of SIX13 (right panel). Kex2 protease was added to the protein at a 1:200 protease to protein ratio prior to crystal tray setup. **(E)** Coomassie-stained gel showing SIX8_C58S^19-141^ (left panel) and mature SIX8_C58S^50-141^ cleaved *in vitro* with thrombin (middle panel). Schematic of SIX8 engineered with an internal thrombin cleavage site replacing the Kex2 cleavage motif (top right panel). Optimised crystals of SIX8_C58S^50-141^ (bottom right panel). **(F)** Coomassie-stained gel showing PSL1_C37S^18-111^ protein (left panel). Optimised crystals of PSL1_C37S^18-111^ (right panel).

**S2 Fig.**
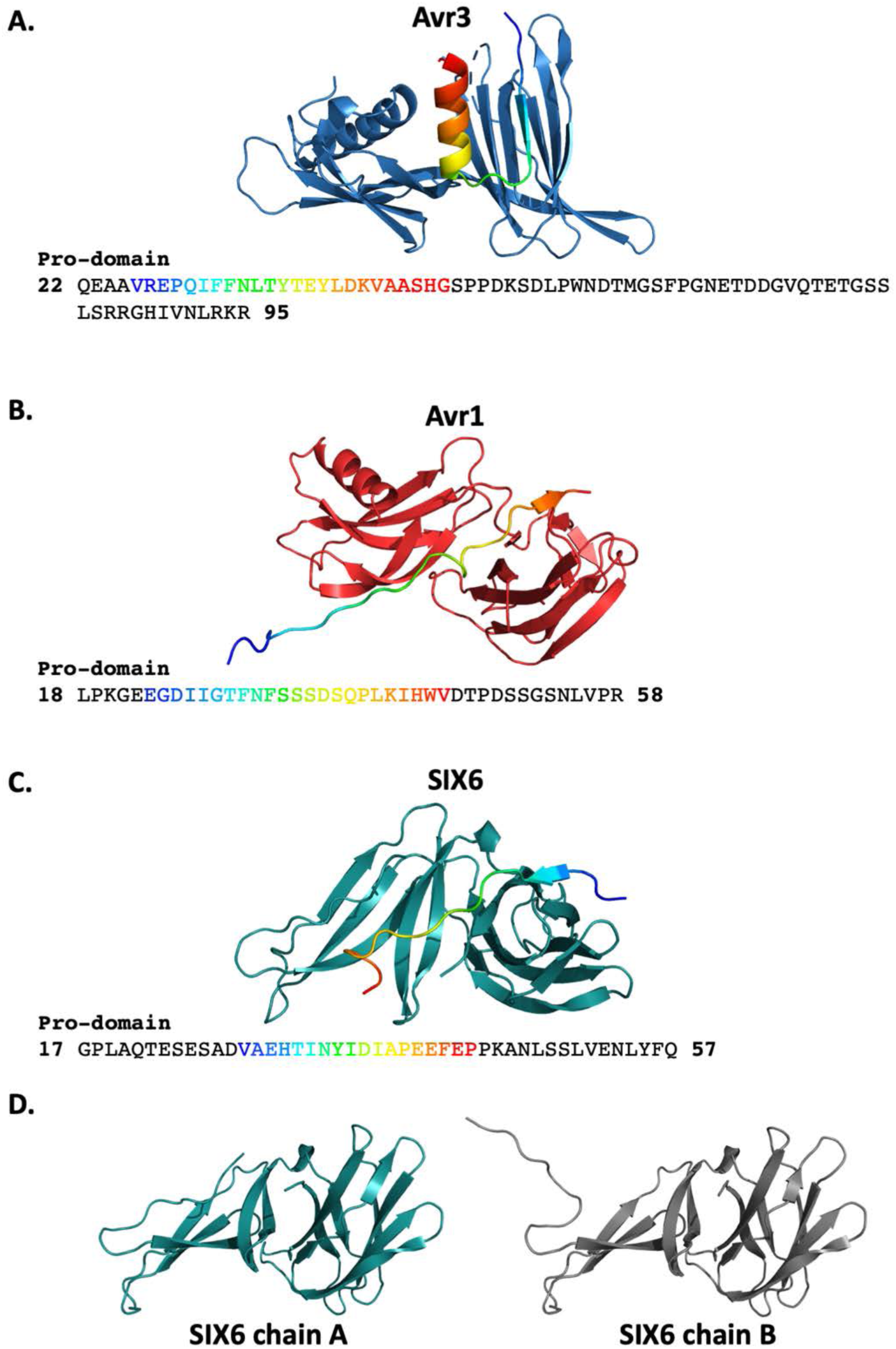
Continuous electron density of the pro-domain is present in the crystal structures of Avr1, Avr3 and SIX6. The crystal structure of **(A)** Avr3, **(B)** Avr1 and **(C)** SIX6 with the pro-domain shown in rainbow (top panels). The amino acid sequence of the pro-domain of Avr3, Avr1 and SIX6 with residues observed in the electron density shown in rainbow text (bottom panels). Residues with no density observed are shown in black. For SIX6, electron density corresponding to the pro-domain was only associated to chain A. **(D)** Different orientations of the N-terminal region of SIX6 between chains A and B. Chain A was used in subsequent structural analysis.

**S3 Fig.**
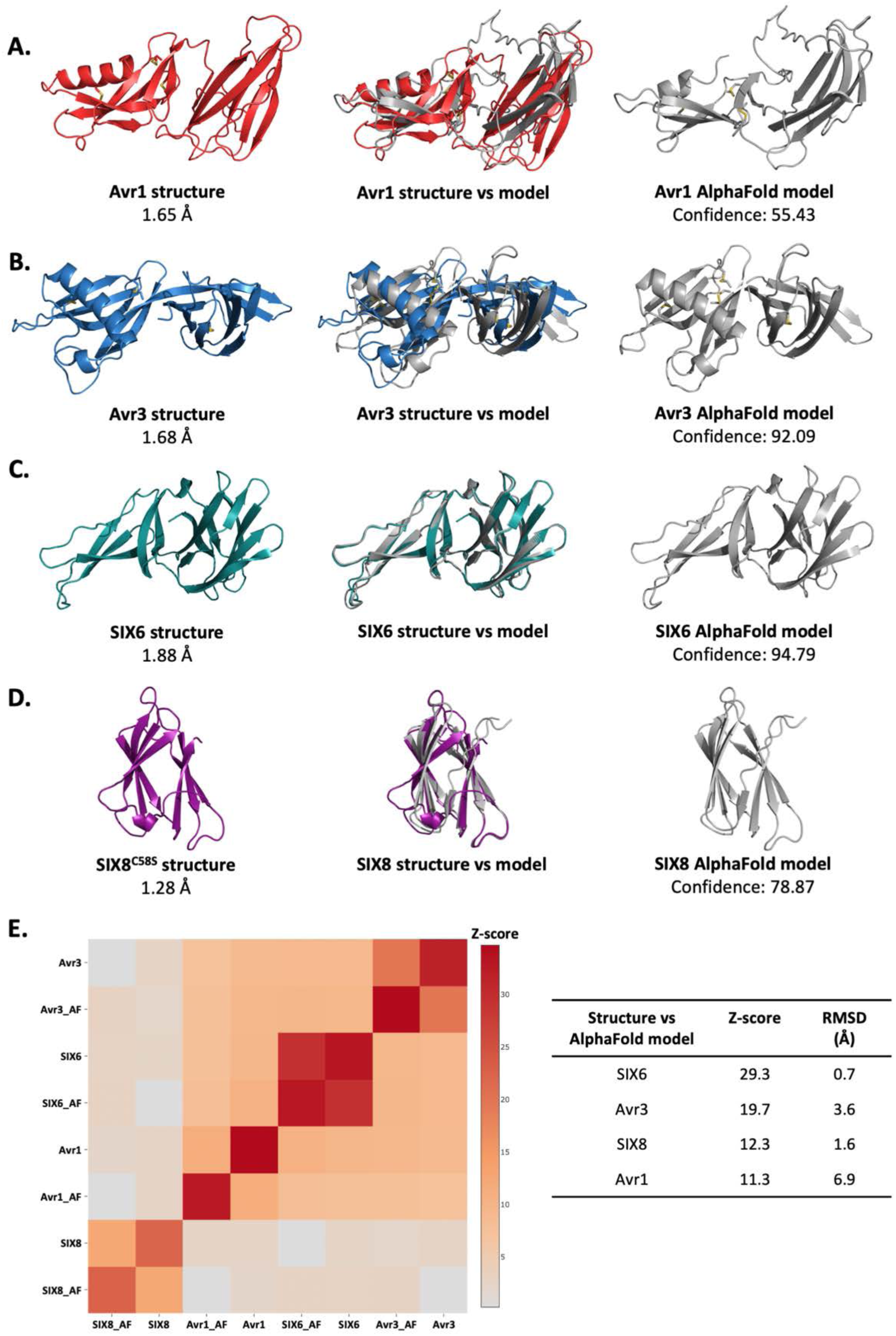
Comparison of AlphaFold2 models against the experimentally solved structures of Avr1, Avr3, SIX6 and SIX8. The crystal structures of (A) Avr1, (B) Avr3, (C) SIX6 and (D) SIX8 (left panels) and AlphaFold2 models [34] (right panels). Crystal structures and AlphaFold2 models of the full structures (middle panels) were superimposed using the pairwise and all against all functions on the DALI server [33]. (E) Heat map of the structural similarity between crystal structures and AlphaFold2 models (left panel). Z-score and RMSD values are shown in the right panel.

**S4 Fig.**
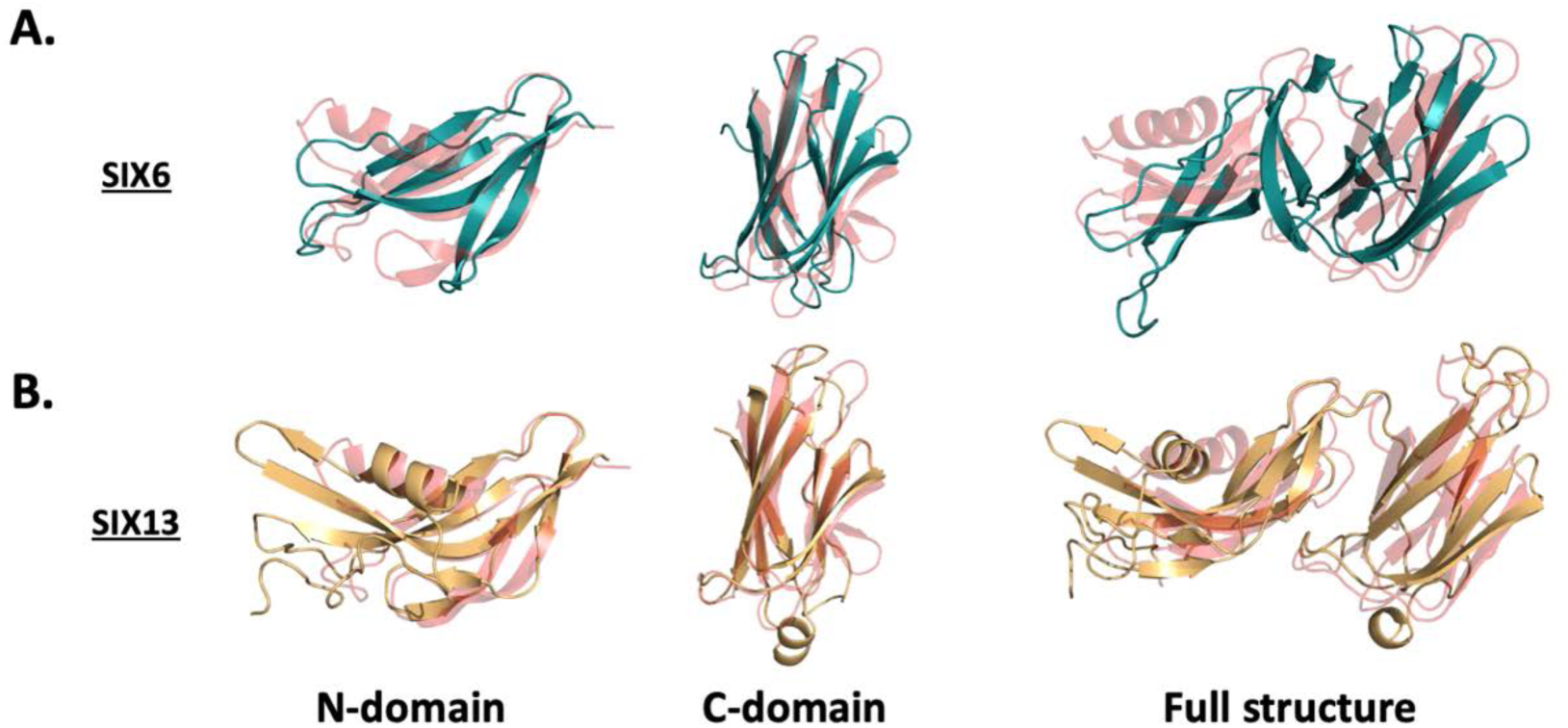
Structural alignments of SIX6 and SIX13 with Avr1. **(A)** SIX6 crystal structure and **(B)** SIX13 AlphaFold2 model aligned with Avr1 using the N-domains alone (left panel), C-domains alone (middle panel) and full structure (right panel). Structural alignment was performed using the pairwise alignment function on the DALI server [33].

**S5 Fig.**
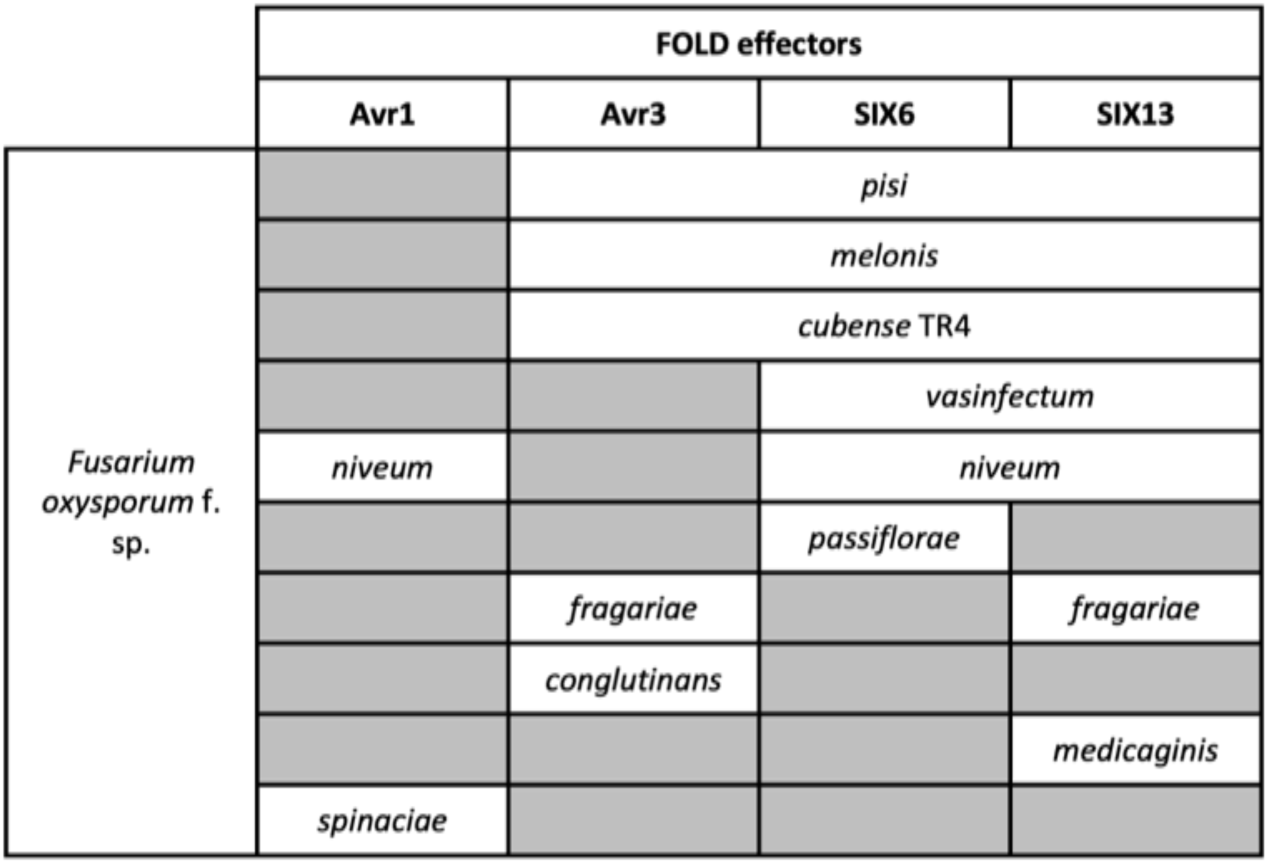
Homologues of FOLD effectors are dispersed across multiple *formae speciales* of *F. oxysporum.* Functional homologues of Avr1 (SIX4), Avr3 (SIX1), SIX6 and SIX13 reported in literature were assessed [6, 8, 35-38].

**S6 Fig.**
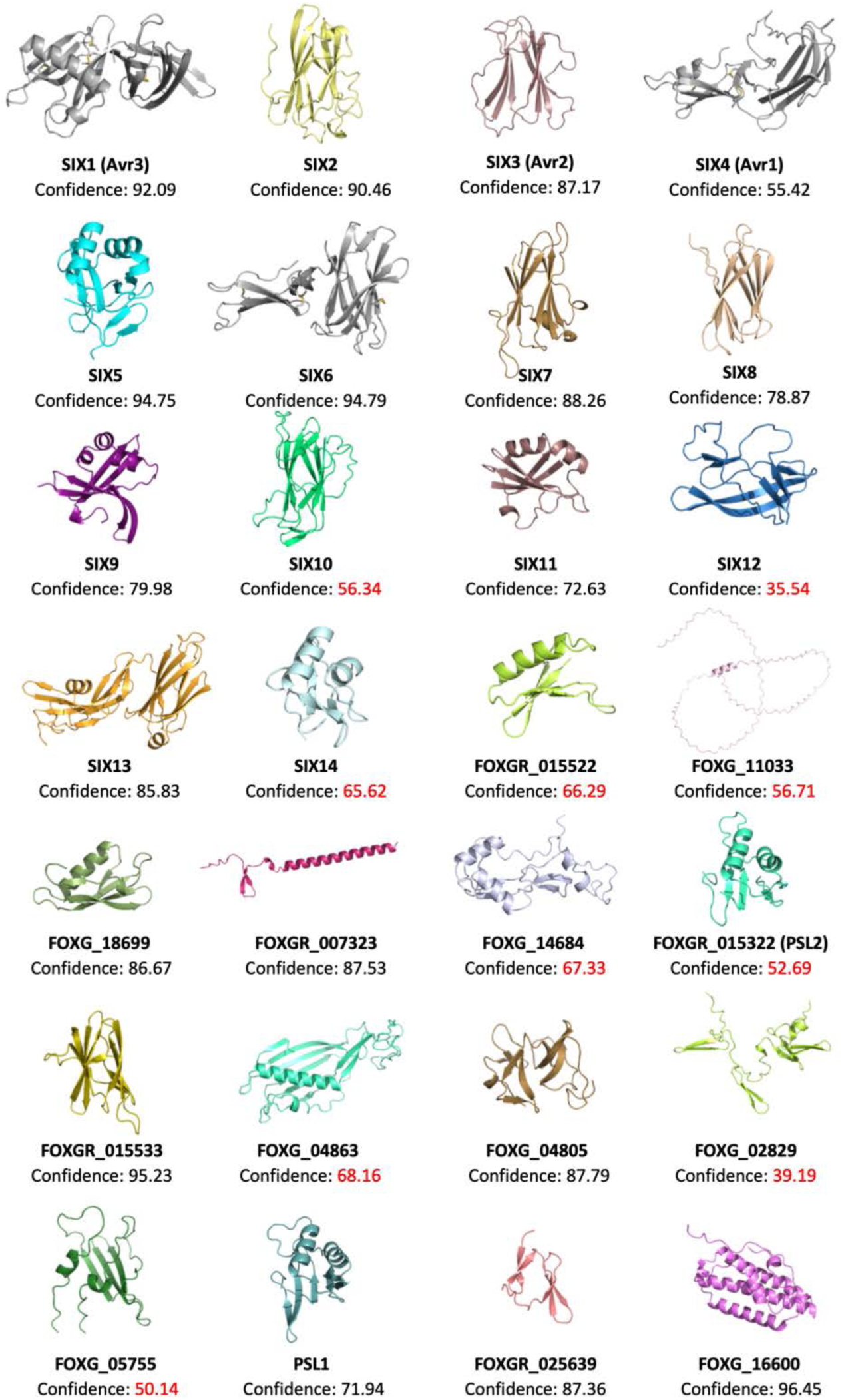
AlphaFold2 models of all SIX effectors and effector candidates. Signal peptides were identified using SignalP-5.0 [92] and removed prior to amino acid sequences being input into AlphaFold2 [34]. Any putative pro-domains were identified by searching for a Kex2-like protease site [31] and removed. The sequence inputs used can be found in S3 Table.

**S7 Fig.**
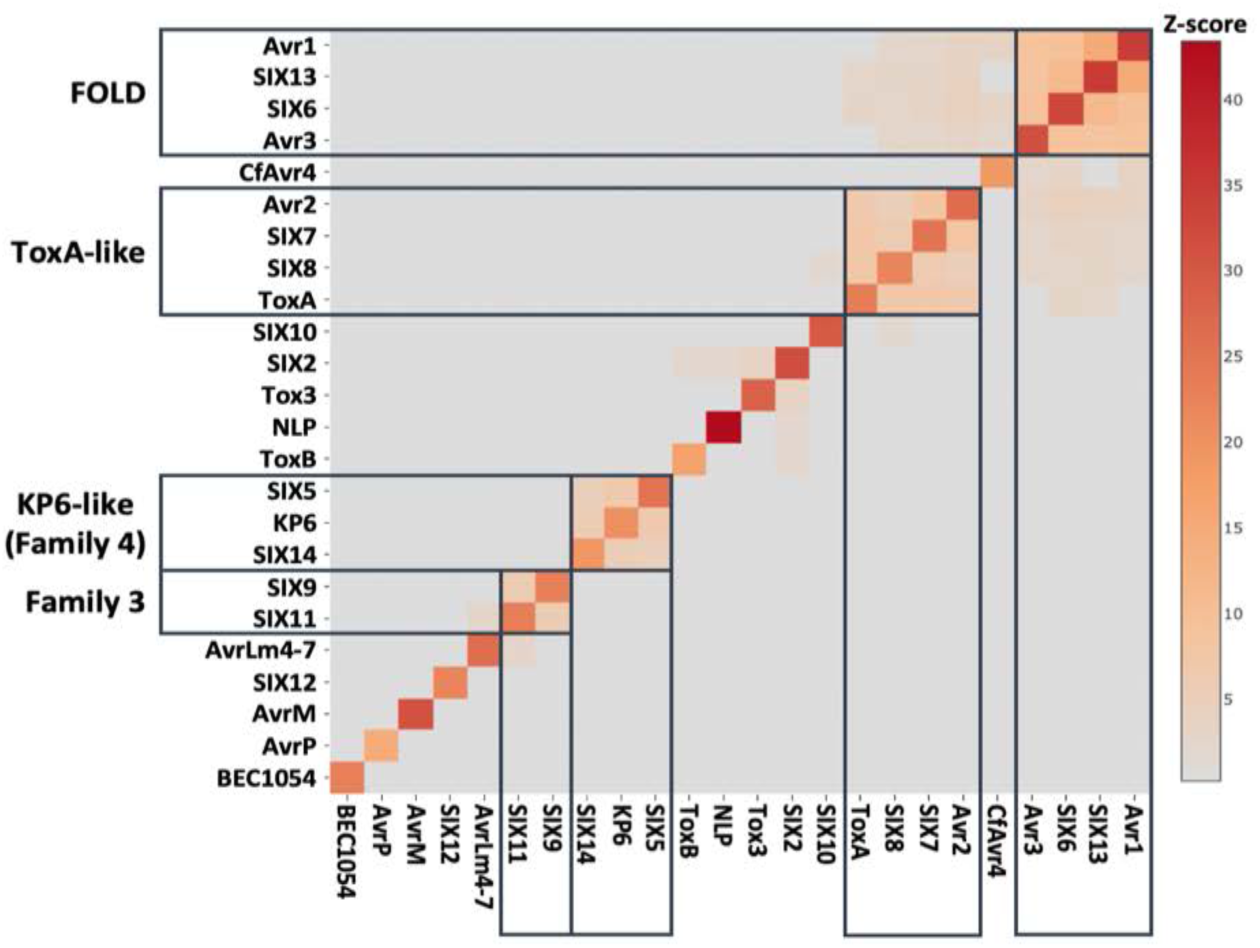
Structural similarity of SIX effectors against representative solved effector structures from known structural families. The solved structures of Avr1, Avr2, Avr3 and SIX6, and AlphaFold2 models the remaining SIX effectors were compared with the structures of ToxA (ToxA-like), ToxB (MAX), Tox3 (Tox3-like), BEC1054 (RALPH), AvrLm4-7 (LARS), AvrP (Zinc finger), CfAvr4 (CBM14-like), AvrM (WY-like), NLP (Actinoporin-like) and KP6 (KP6-like). Structural alignment was performed using the all against all function on the DALI server [33]. Structural similarity was measured using Z-score. Groupings with Z-scores > 2 were outlined.

**S8 Fig.**
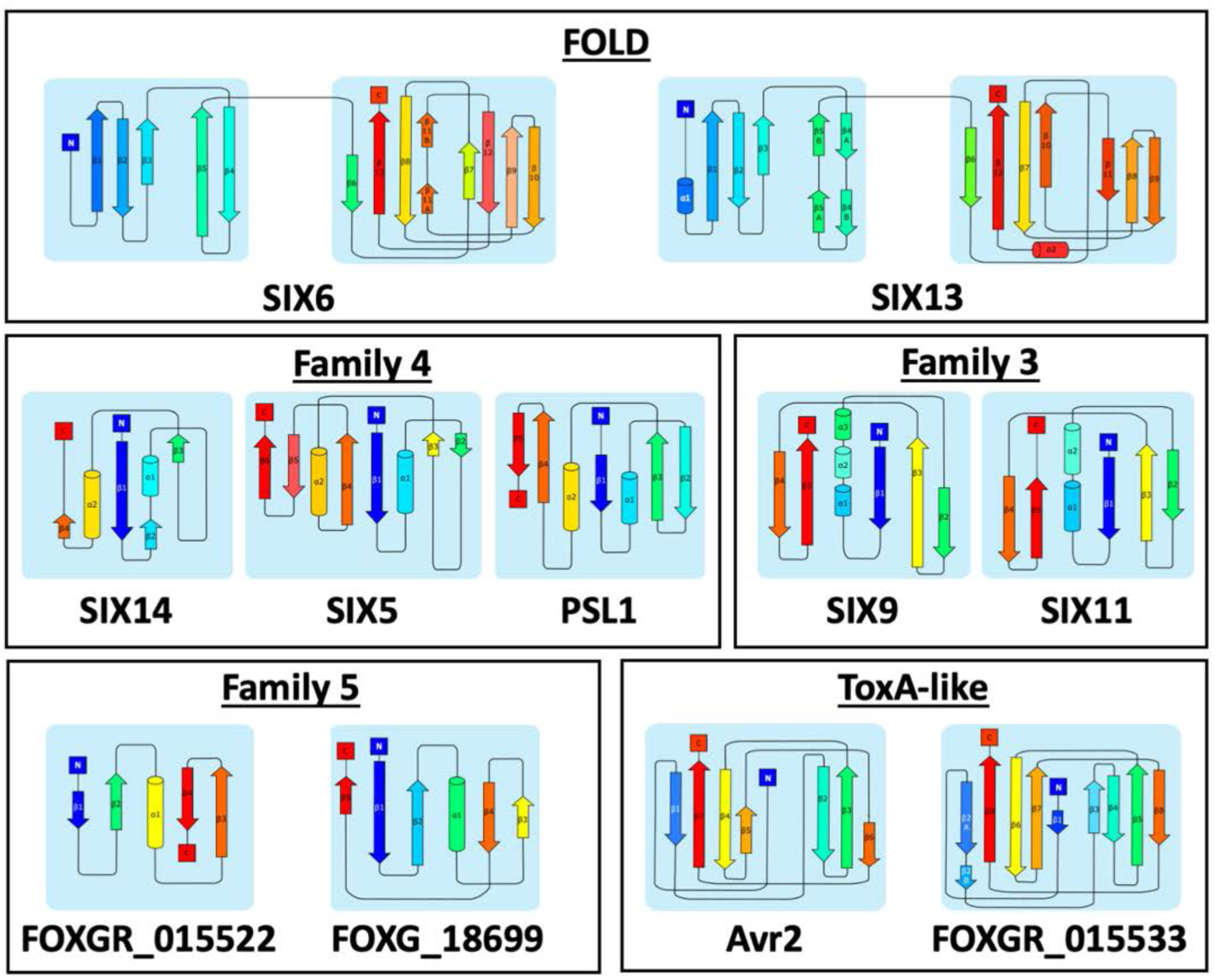
Secondary structure topology maps of representative SIX structural family members. Secondary topology maps of AlphaFold2 models were generated using Pro-origami [94] and manually edited in Inkscape. The β-strands and α-helices are represented by arrows and cylinders, respectively. The secondary structural elements are coloured in rainbow, from blue at the N-terminus to red at the C-terminus.

**S9 Fig.**
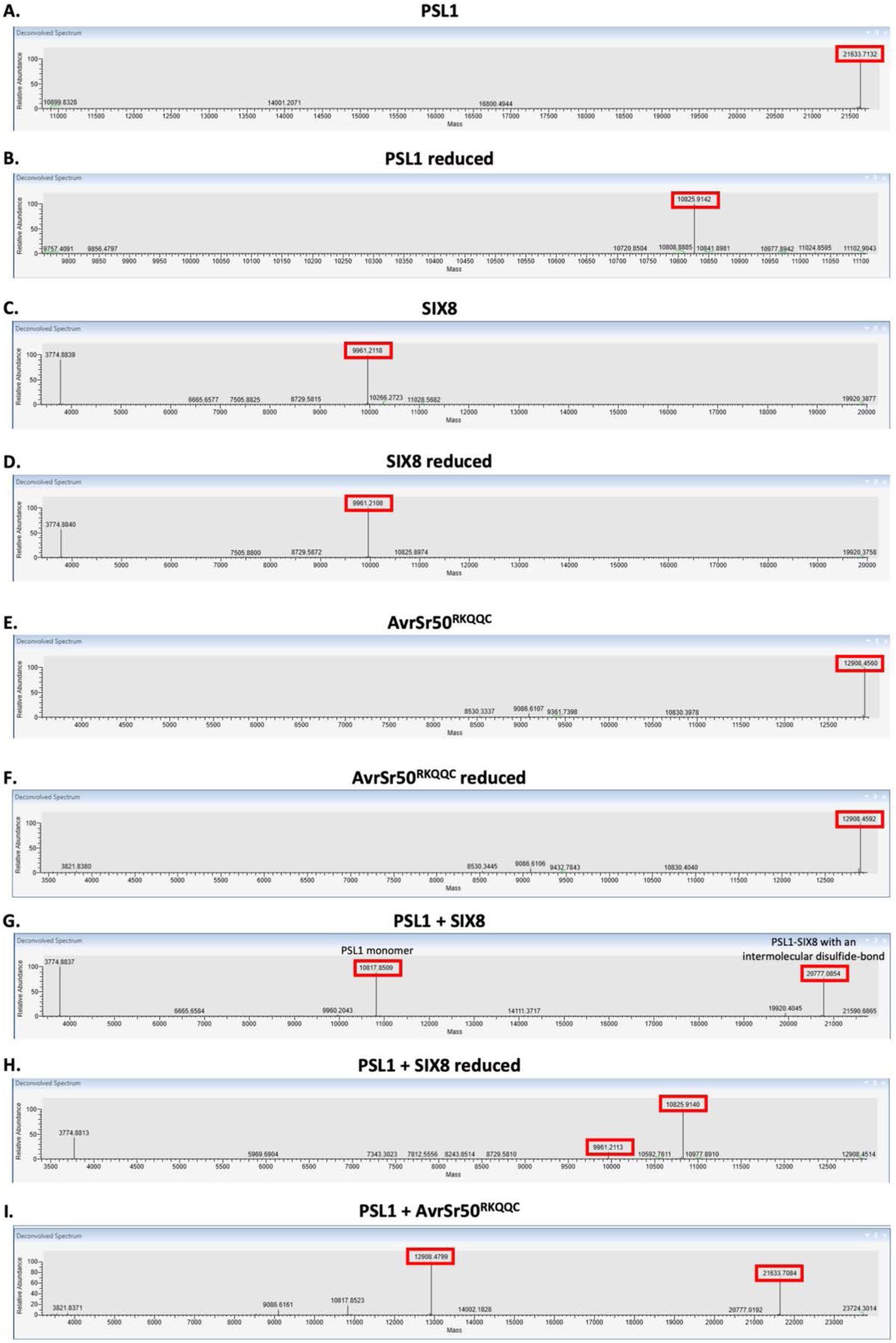
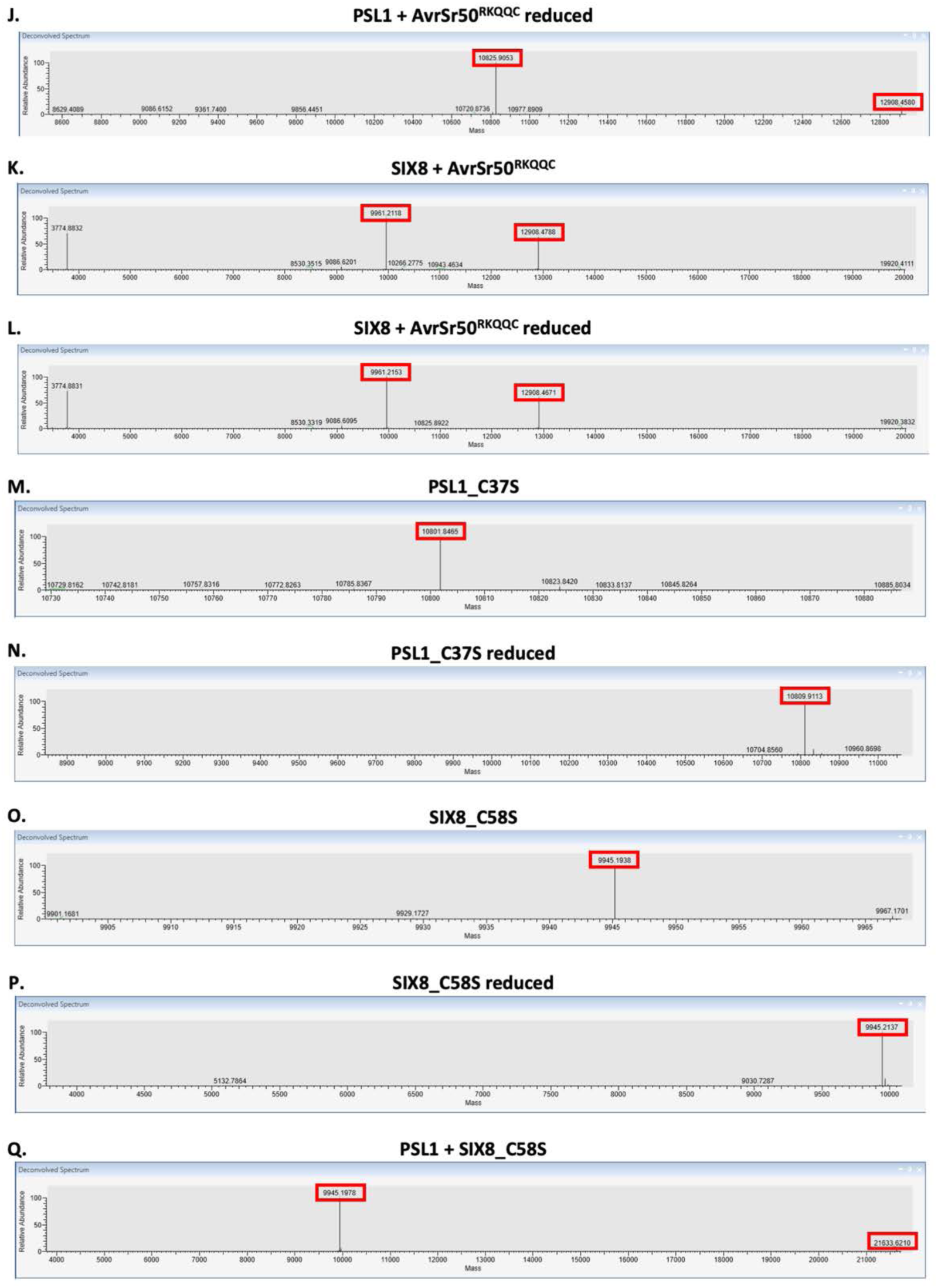
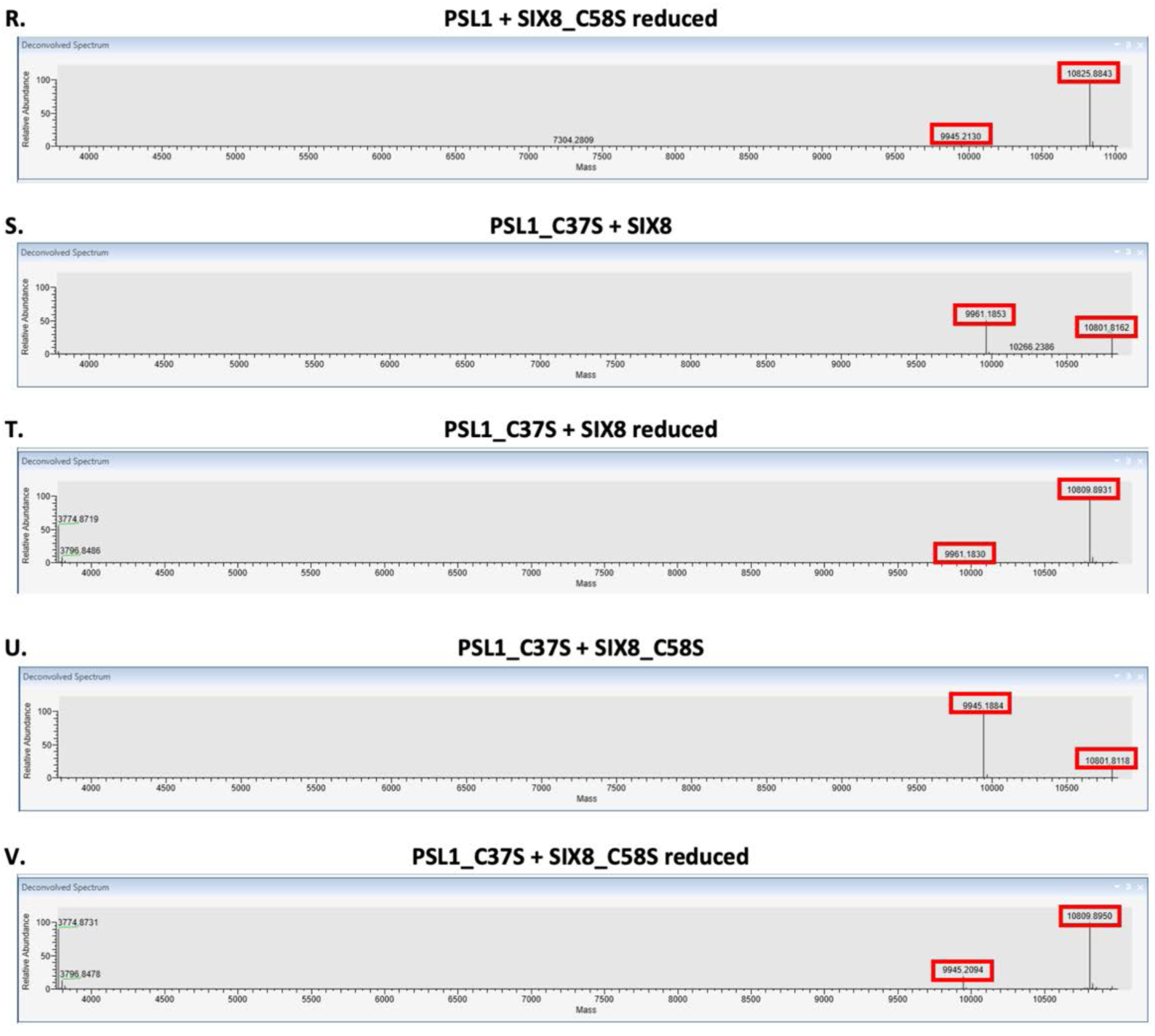
Intact mass spectrometry analysis of the PSL1-SIX8 interaction. Deconvoluted mass spectra of **(A)** PSL1, **(B)** reduced PSL1, **(C)** SIX8, **(D)** reduced SIX8, **(E)** AvrSr50^RKQQC^, **(F)** reduced AvrSr50^RKQQC^, **(G)** PSL1 + SIX8, **(H)** reduced PSL1 + SIX8, **(I)** PSL1 + AvrSr50^RKQQC^, **(J)** reduced PSL1 + AvrSr50^RKQQC^, **(K)** SIX8 + AvrSr50^RKQQC^, **(L)** reduced SIX8 + AvrSr50^RKQQC^, **(M)** PSL1_C37S, **(N)** reduced PSL1_C37S, **(O)** SIX8_C58S, **(P)** reduced SIX8_C58S, **(Q)** PSL1 + SIX8_C58S, **(R)** reduced PSL1 + SIX8_C58S, **(S)** PSL1_C37S + SIX8, **(T)** reduced PSL1_C37S + SIX8, **(U)** PSL1_C37S + SIX8_C58S, **(V)** reduced PSL1_C37S + SIX8_C58S. Reduced samples were heated with DTT prior to running the samples.

**S10 Fig.**
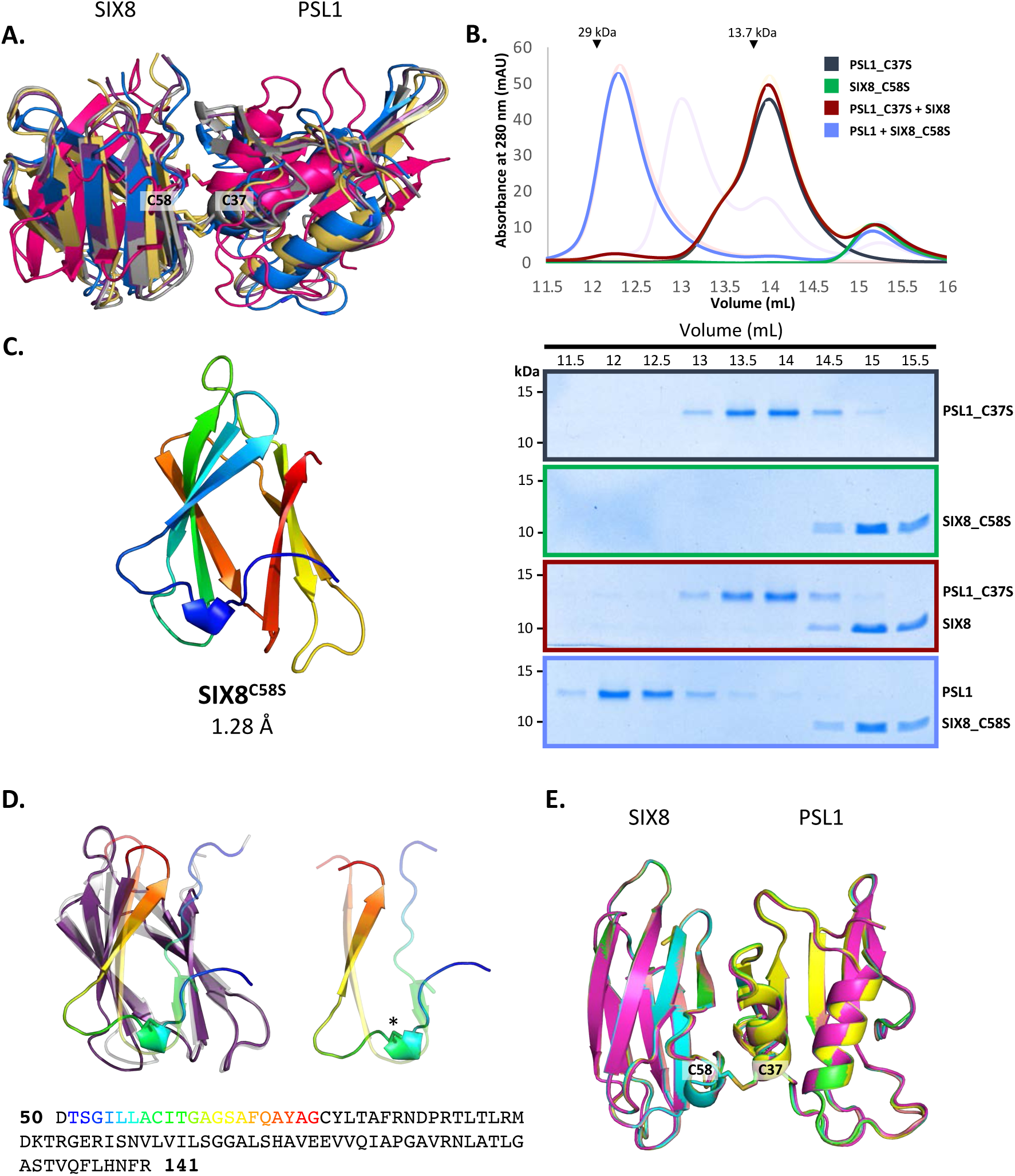
Interaction between PSL1 and SIX8 mutants. **(A)** Model of the SIX8-PSL1 complex generated by AlphaFold2-Multimer (five models shown). Co-localisation of Cys 58 from SIX8 and Cys 37 from PSL1 shown in stick. **(B)** Size exclusion chromatograms of PSL1_C37S alone (black), SIX8_C58S alone (green), PSL1_C37S and SIX8 (maroon), and PSL1 and SIX8_C58S (light purple) following a 30 min incubation separated on a Superdex S75 Increase 10/300 SEC column (top panel). Equal concentrations of the protein were used (note the absorbance of SIX8 at 280 nm is ∼0.3 resulting in a smaller absorbance and peak height). Indicated sizes above the chromatogram are based on protein standards run under similar conditions as presented in the manufacturer’s column guidelines. Transparent peaks correspond to the size exclusion chromatograms of PSL1 alone (red), SIX8 alone (blue), PSL1 and SIX8 (purple) found in Figure 4B. Coomassie-stained SDS-PAGE gels depicting samples taken from 500 µL fractions corresponding to the volumes indicated above the gels, with molecular weights (left) and proteins (weight) annotated (bottom panel). **(C)** Cartoon representation of the crystal structure of SIX8^C58S^ at 1.28 Å resolution, coloured from N (blue) to C (red) terminus. **(D)** Comparison of the SIX8 structure and the AlphaFold2 model. The SIX8 structure (purple) and AlphaFold2 model (grey) were superimposed using the DALI server (top panel) [33]. The N-terminus is coloured in rainbow. The location of C58S is shown by an asterisk. Amino acid sequence of SIX8 with residues of the N-terminus in rainbow corresponding to the structure (bottom panel). **(E)** Model of the SIX8-PSL1 complex generated by AlphaFold2-Multimer (five models shown), when the SIX8^C58S^ structure was used as a template. Co-localisation of Cys 58 from SIX8 and Cys 37 from PSL1 shown in stick.

**S11 Fig.**
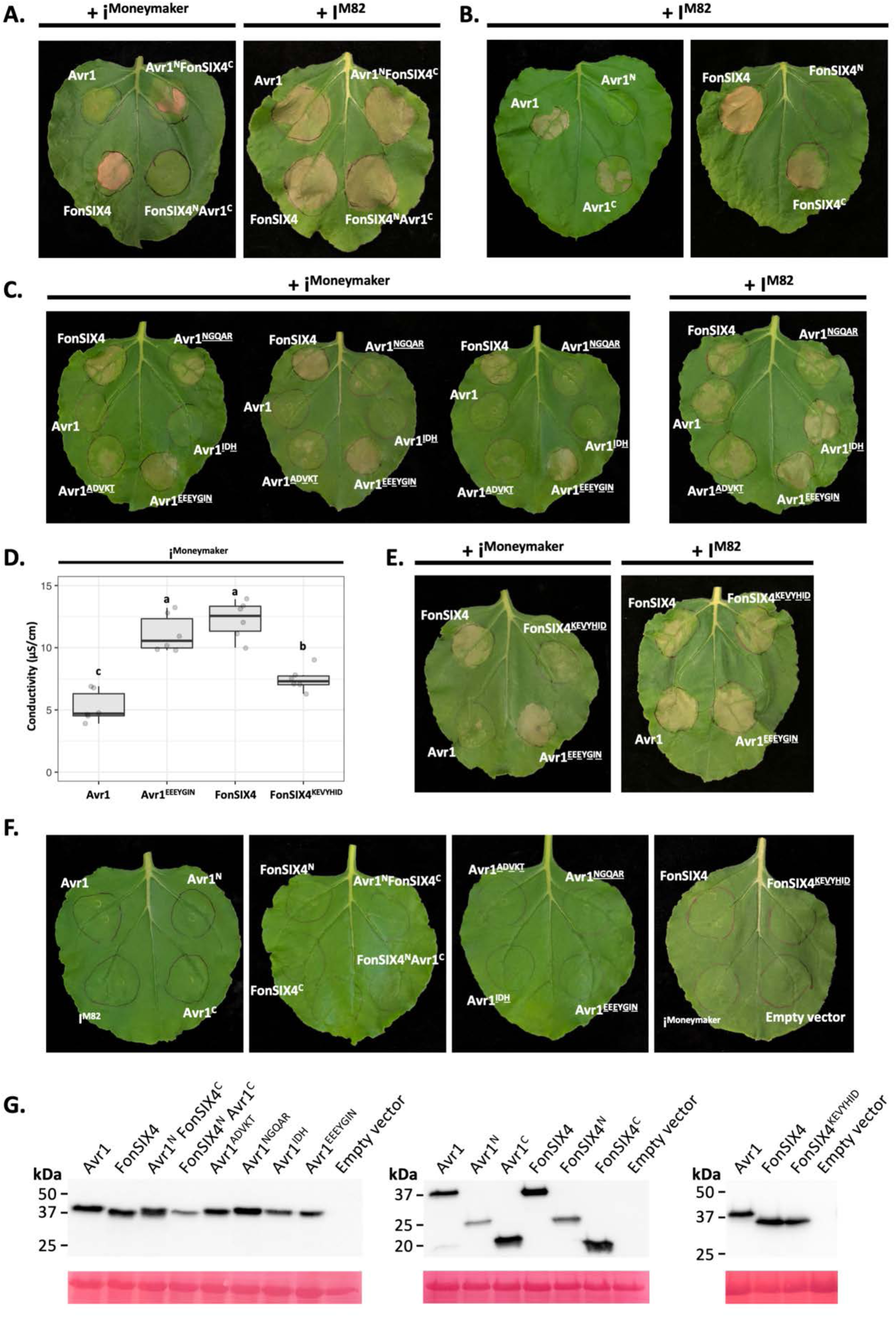
**(A)** i^Moneymaker^ recognises FonSIX4 at the C-domain. N and C-domain swapped chimeras of Avr1 and FonSIX4 were expressed with i^Moneymaker^ (left panel) and I^M82^ (right panel). **(B)** The C-domains of Avr1 and FonSIX4 are sufficient for I^M82^ recognition. N and C-domains of Avr1 (left panel) and FonSIX4 (right panel) were expressed with I^M82^. **(C)** Variation in the recognition response of Avr1 mutants by i^Moneymaker^. Avr1 mutants (Avr1<u>^A^</u>^D<u>V</u>KT^, Avr1<u>^NGQAR^</u>, Avr1^ID<u>H</u>^, Avr1^EEEY<u>GIN</u>^) were co-expressed with i^Moneymaker^ (left panel) or I^M82^ (right panel). **(D)** Ion leakage quantification of the reciprocal Avr1^EEEY<u>GIN</u>^ and FonSIX4^KE<u>V</u>Y<u>HID</u>^ mutants when transiently co-expressed with i^Moneymaker^. **(E)** Leaf images of Avr1^EEEY<u>GIN</u>^ and FonSIX4^KE<u>V</u>Y<u>HID</u>^ mutants transiently co-expressed with i^Moneymaker^ (left panel) or I^M82^ (right panel). Polymorphic residues between Avr1 and FonSIX4 are underlined. **(F)** Effectors and receptors expressed alone do not cause cell death in *N. benthamiana.* All Avr1, FonSIX4 effectors and I receptors were transiently expressed in *N. benthamiana* via *Agrobacterium*-mediated transformation. Six biological replicates were assessed for all qualitative assessment of I receptor mediated cell death in *N. benthamiana*. All leaves were imaged 4 – 7 dpi. **(G)** Western blots of Avr1 and FonSIX4 constructs with a C-terminal HA tag. Total proteins were extracted from *N. benthamiana* leaves 3 dpi and separated by SDS-PAGE. The samples were transferred onto a membrane, probed anti-HA antibodies and analysed under a chemiluminescence imager. The membrane was stained with Ponceau S to show equal sample loading.

**S12 Fig.**
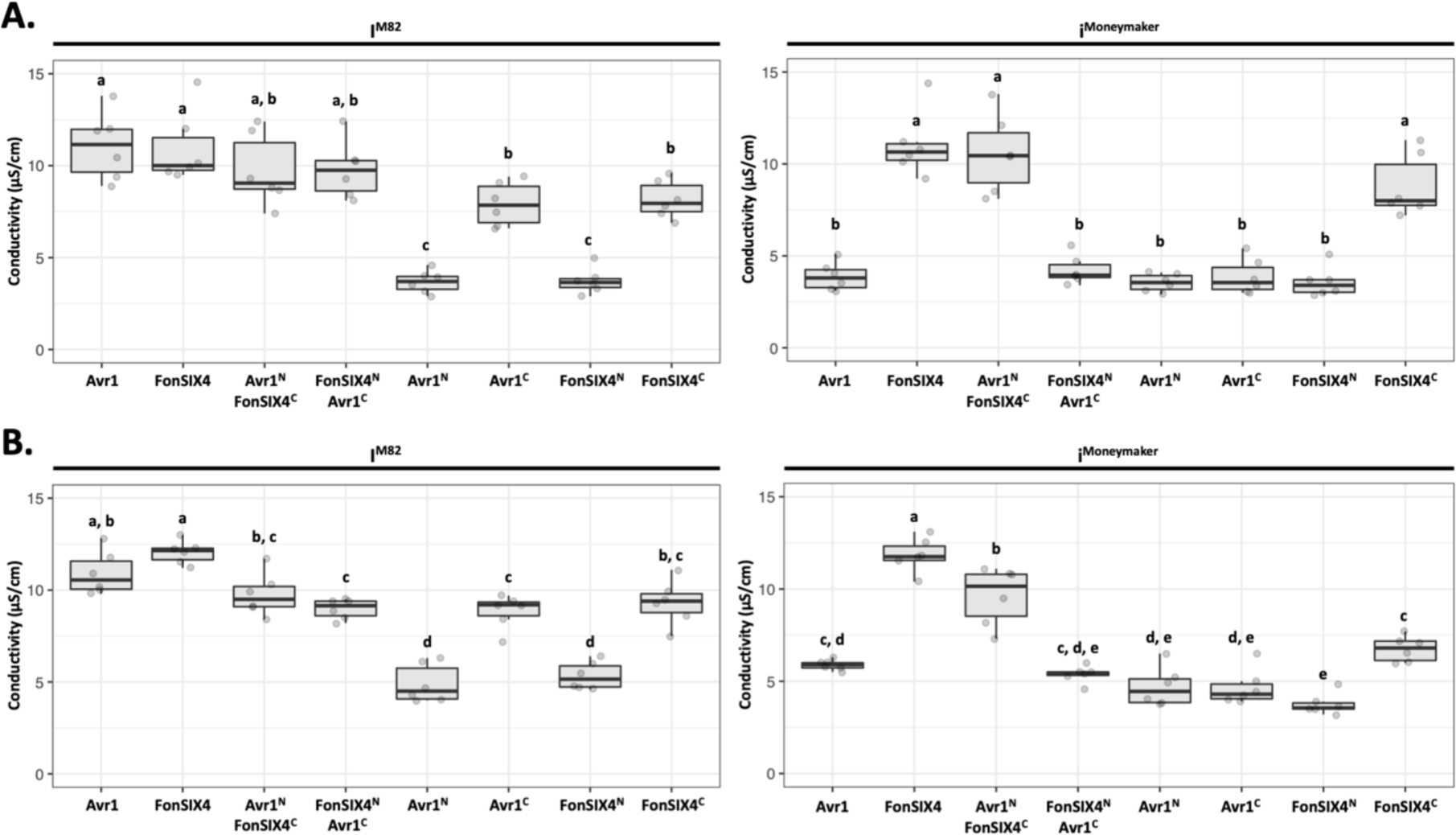
A & B. Independent ion leakage experiments of different Avr1 and FonSIX4 constructs expressed with I^M82^ or i^Moneymaker^. Avr1 and FonSIX4 effector constructs were transiently expressed with I^M82^ (left panels) or i^Moneymaker^ (right panels) via *Agrobacterium-*mediated transformation in *N. benthamiana*. Six biological replicates each consisting of three leaf discs per leaf were harvested 24 hours post infiltration and incubated in water for 30 minutes. The water was replaced and the conductivity was measured after 24-48 hours. One-way ANOVA and post *hoc* Tukey’s honestly significant difference tests were performed. Treatments that do not share a letter are significantly different from each other at *p < 0.05*.

**S13 Fig.**
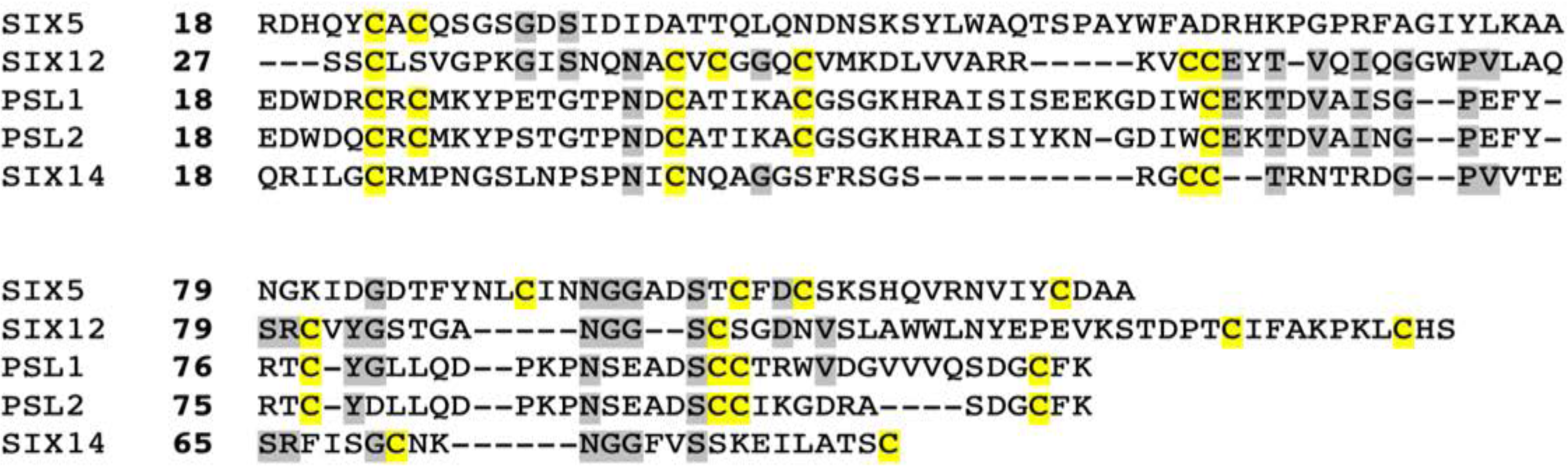
Amino acid sequence alignment of SIX12 against Family 4 members reveals a similar cysteine spacing. All protein sequences have their signal peptides removed. The cysteine residues are highlighted in yellow and groups of two or more amino acid residues shared with SIX12 are highlighted in grey.

**S14 Fig.**
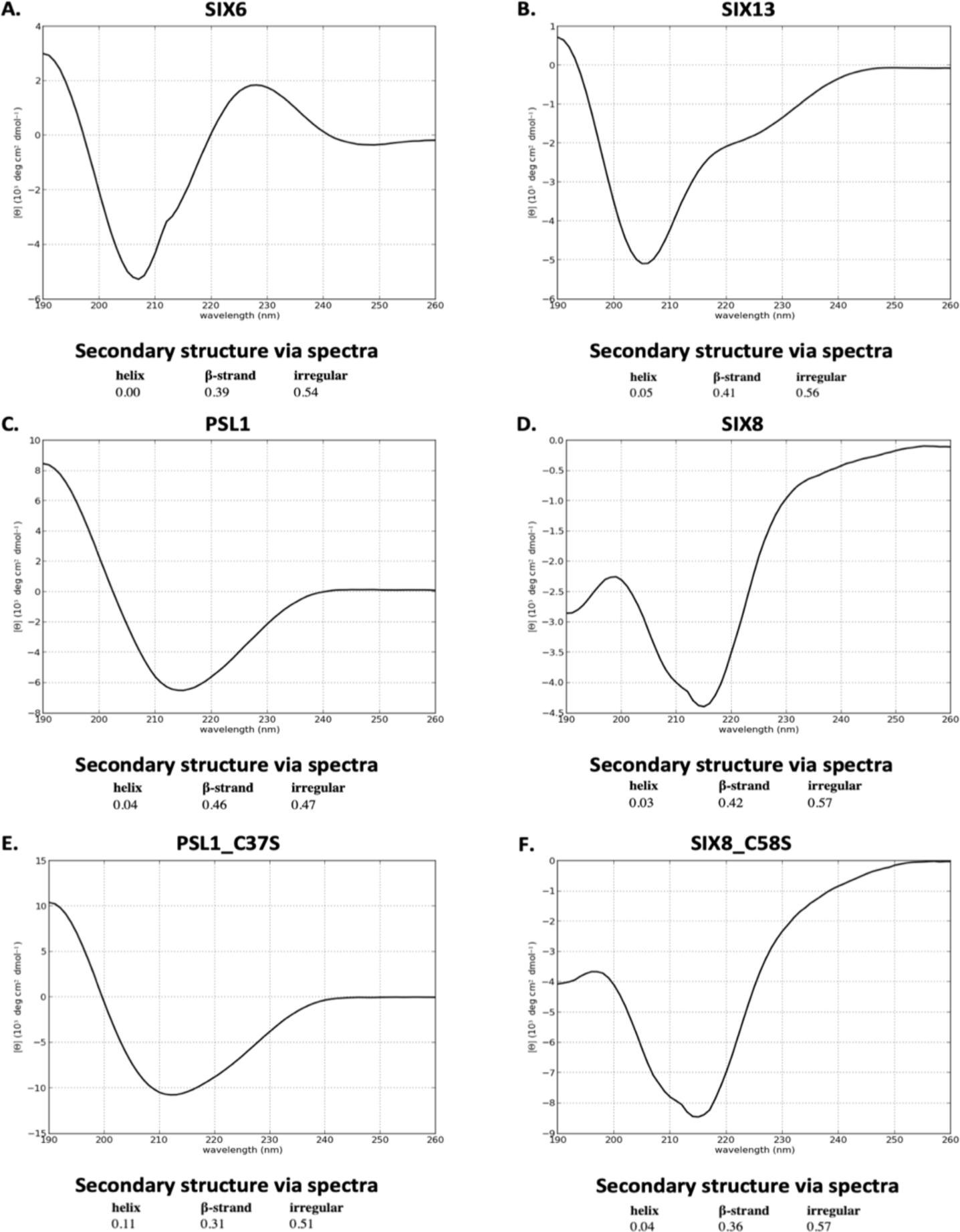
Circular dichroism analysis of purified recombinant proteins. CD spectra of **(A)** SIX6, **(B)** SIX13, **(C)** PSL1, **(D)** SIX8, **(E)** PSL1_C37S and **(F)** SIX8_C58S proteins were plotted, and secondary structure elements analysed using the CAPITO webserver [81].

## Supplementary tables

**S1 Table.**
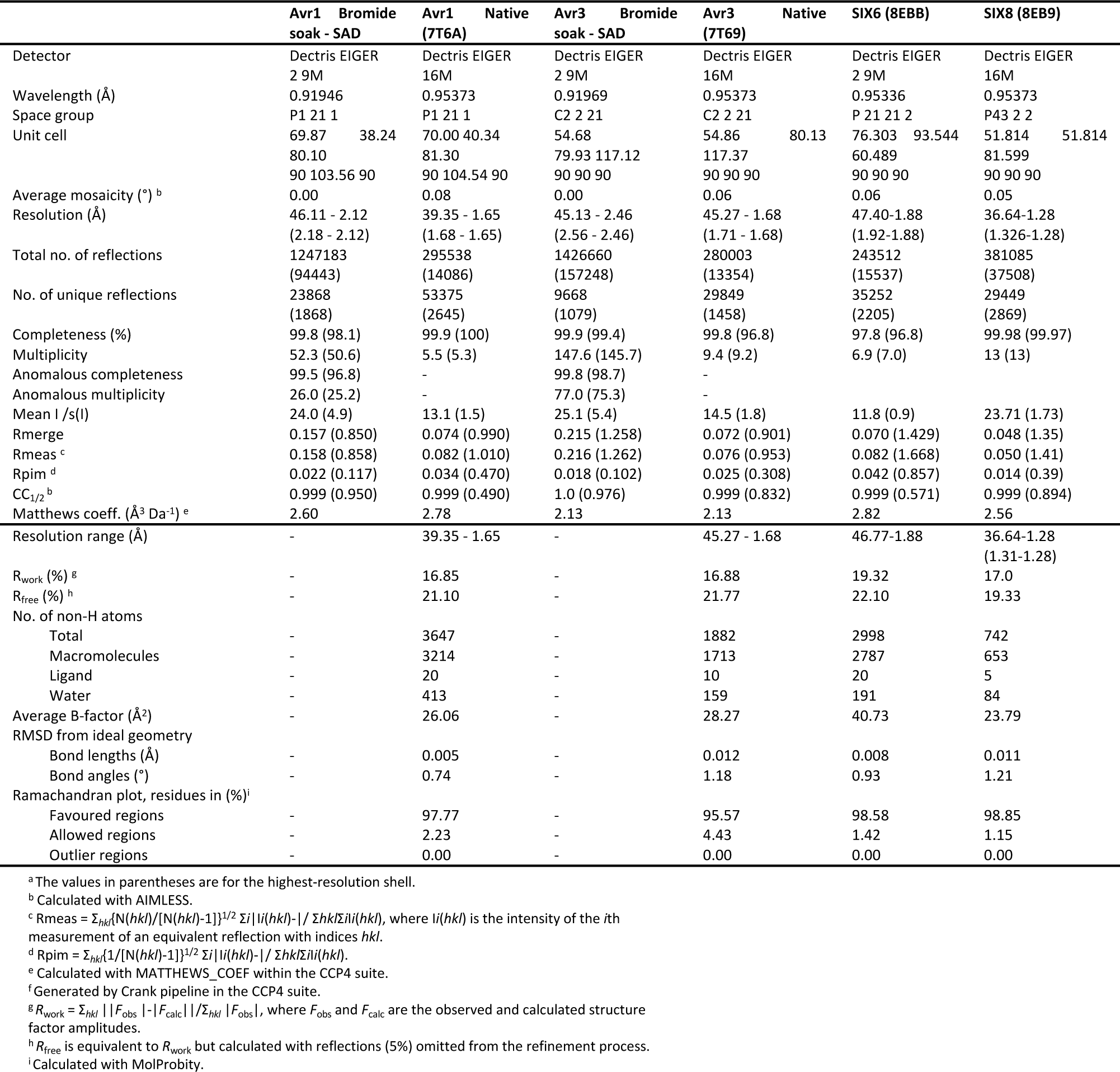
X-ray data collection, structure solution and refinement statistics for Avr1, Avr3, SIX6 and SIX8.

**S2 Table.** (supplementary file)

**S3 Table.**
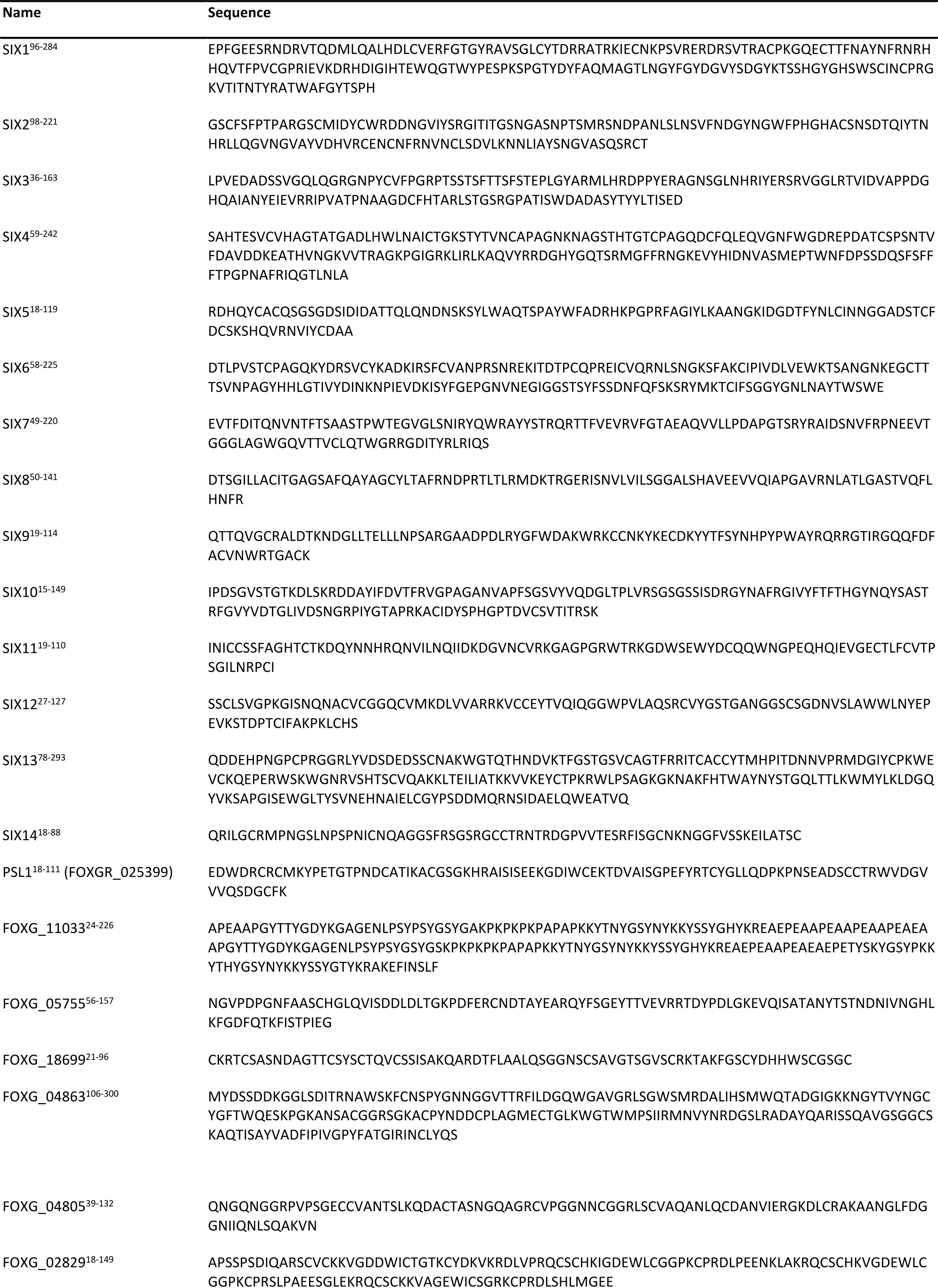
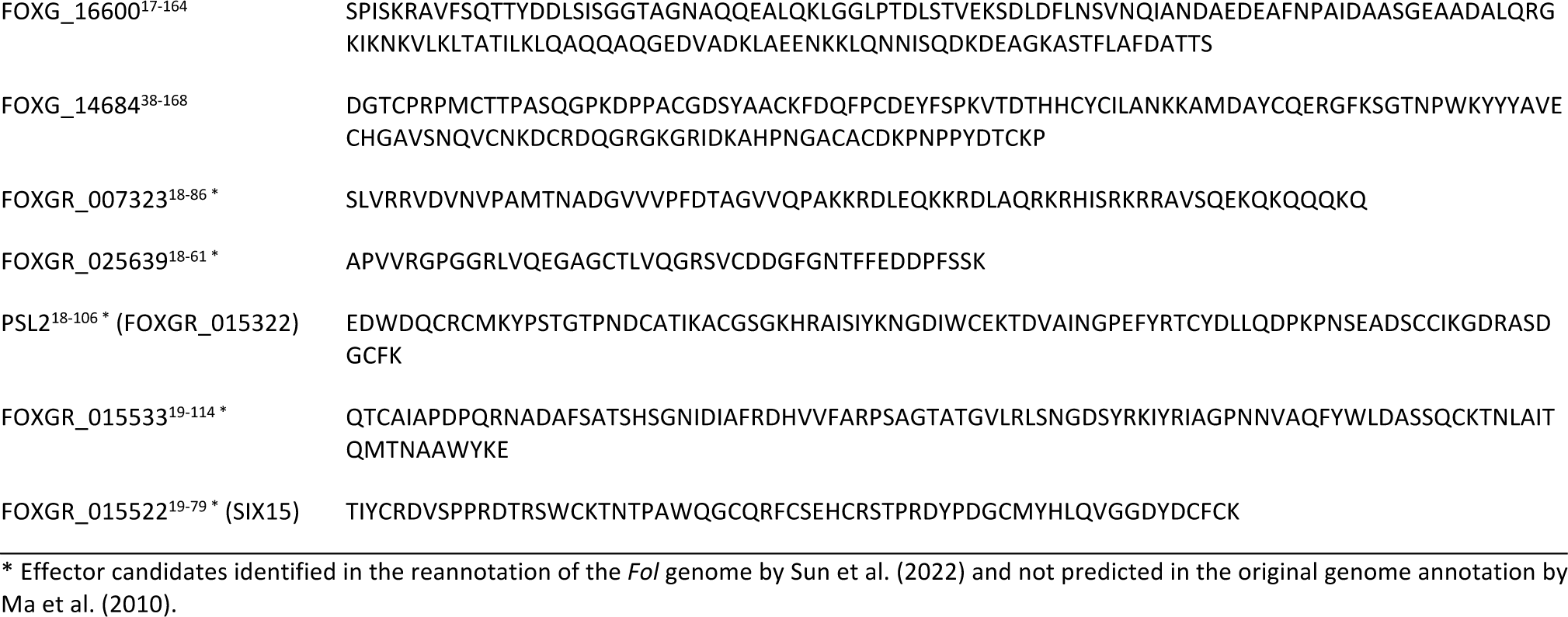
Amino acid sequence inputs for AlphaFold2.

**S4 Table.**
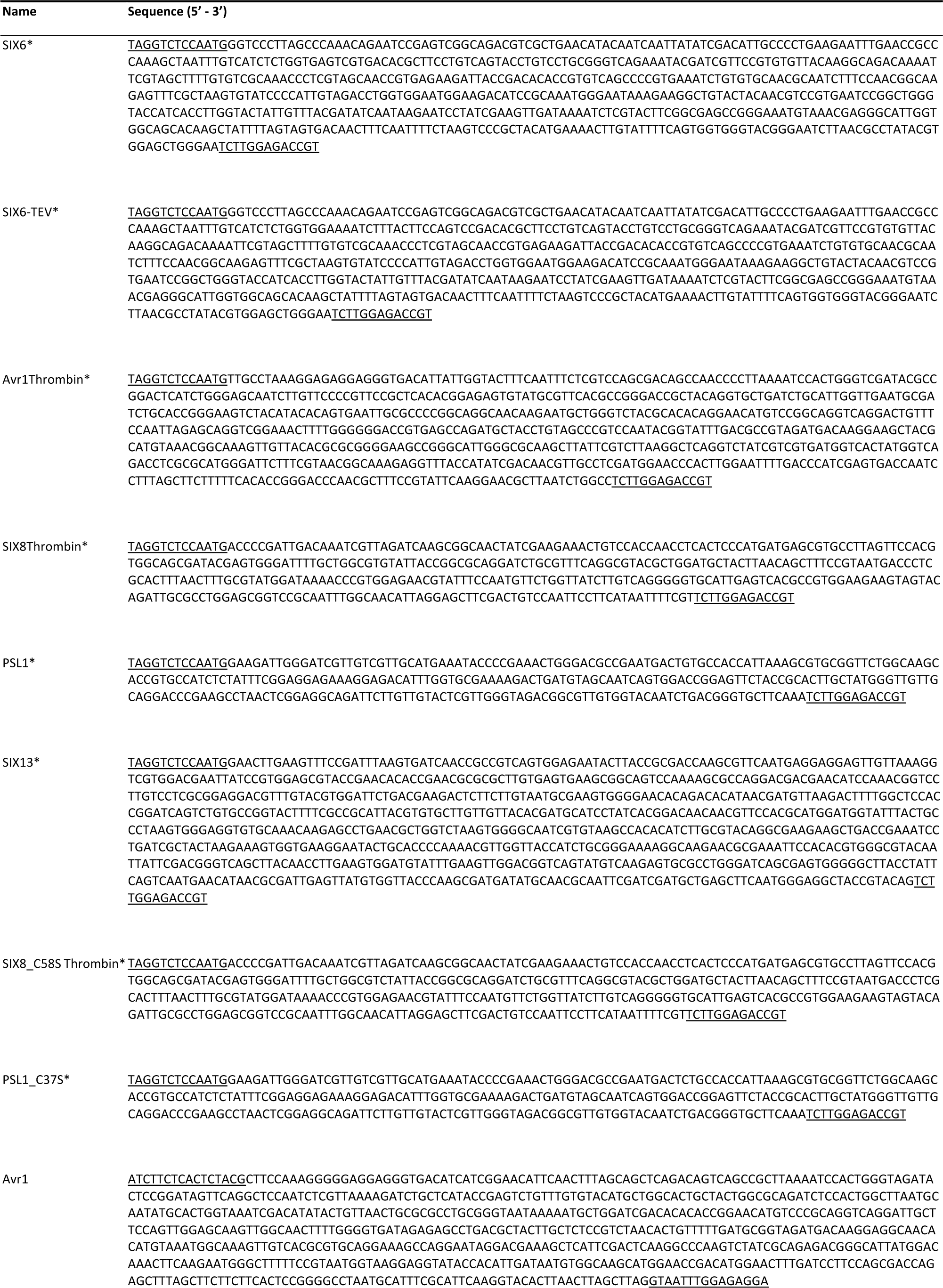
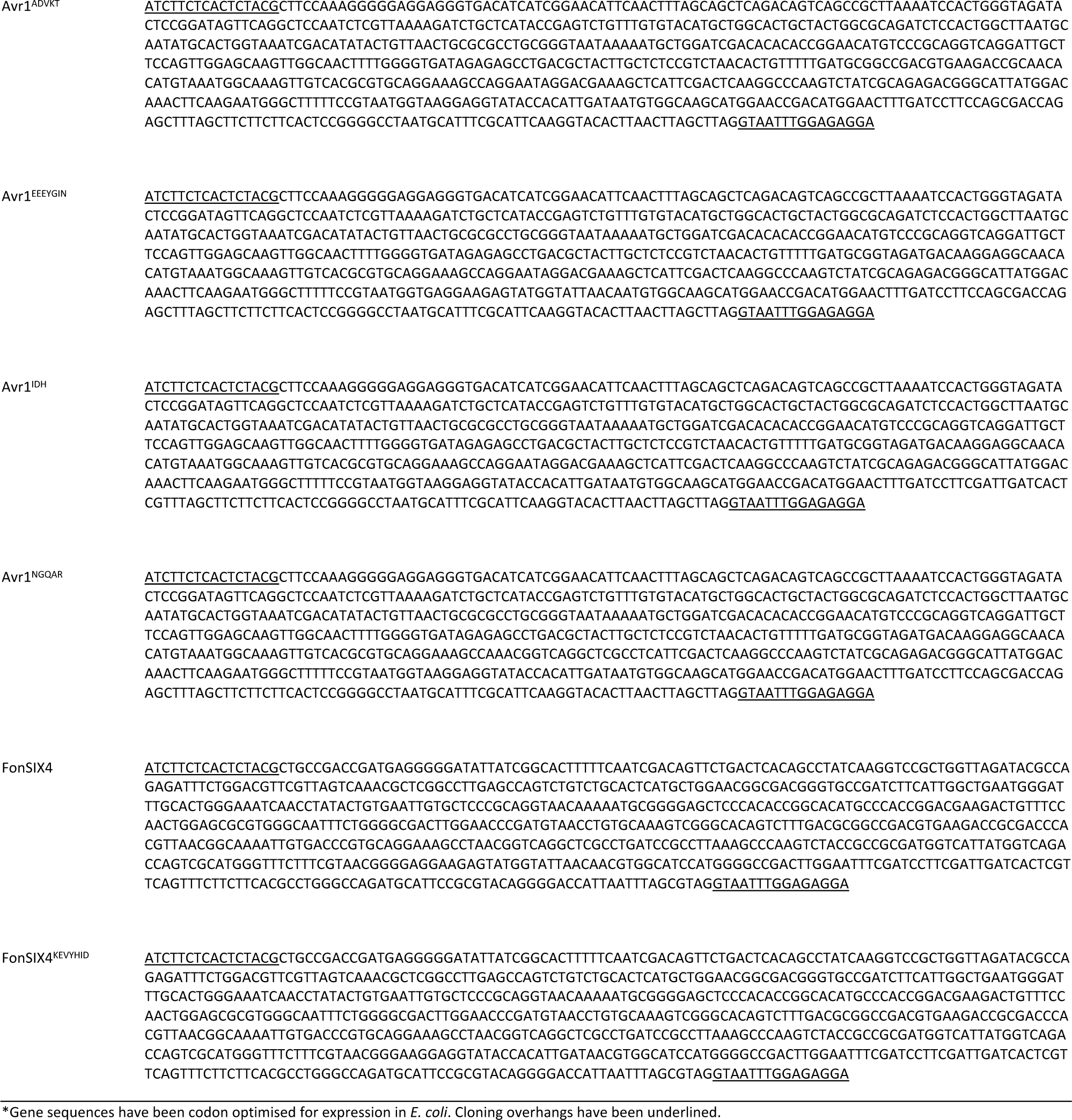
DNA sequences of synthesised gene fragments used in this study.

**S5 Table.**
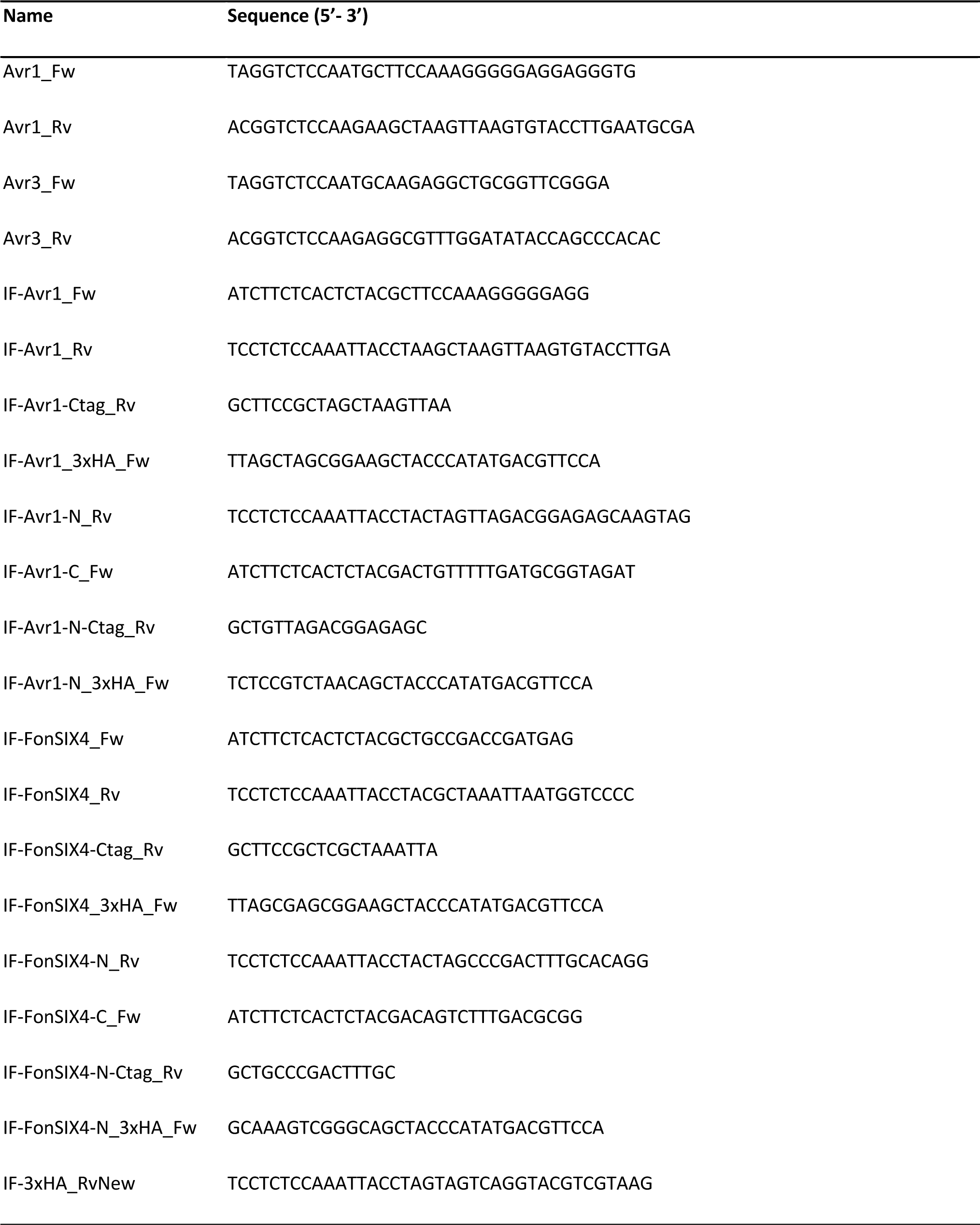
Primers used in this study.

## Literature Cited

1. Dean R, Van Kan JAL, Pretorius ZA, Hammond-Kosack KE, Di Pietro A, Spanu PD, et al. The Top 10 fungal pathogens in molecular plant pathology. Mol Plant Pathol. 2012;13(4):414–430.

2. Rep M. Small proteins of plant-pathogenic fungi secreted during host colonization. FEMS Microbiol Lett. 2005;253(1):19–27.

3. Houterman PM, Speijer D, Dekker HL, de Koster CG, Cornelissen BJC, Rep M. The mixed xylem sap proteome of *Fusarium oxysporum*-infected tomato plants. Mol Plant Pathol. 2007;8(2):215–221.

4. Ma LJ, van der Does HC, Borkovich KA, Coleman JJ, Daboussi MJ, Di Pietro A, et al. Comparative genomics reveals mobile pathogenicity chromosomes in *Fusarium*. Nature. 2010;464(7287):367–373.

5. Rep M, van der Does HC, Meijer M, van Wijk R, Houterman PM, Dekker HL, et al. A small, cysteine-rich protein secreted by *Fusarium oxysporum* during colonization of xylem vessels is required for *I-3*-mediated resistance in tomato. Mol Microbiol. 2004;53(5):1373–1383.

6. Schmidt SM, Houterman PM, Schreiver I, Ma L, Amyotte S, Chellappan B, et al. MITEs in the promoters of effector genes allow prediction of novel virulence genes in *Fusarium oxysporum*. BMC Genomics. 2013;14:119.

7. Vlaardingerbroek I, Beerens B, Rose L, Fokkens L, Cornelissen BJ, Rep M. Exchange of core chromosomes and horizontal transfer of lineage-specific chromosomes in *Fusarium oxysporum*. Environ Microbiol. 2016;18(11):3702–3713.

8. Gawehns F, Houterman PM, Ait Ichou F, Michielse CB, Hijdra M, Cornelissen BJC, et al. The *Fusarium oxysporum* effector Six6 contributes to virulence and suppresses *I-2*-mediated cell death. Mol Plant Microbe Interact. 2014;27(4):336–348.

9. Ma L, Houterman PM, Gawehns F, Cao L, Sillo F, Richter H, et al. The *AVR2-SIX5* gene pair is required to activate *I-2*-mediated immunity in tomato. New Phytol. 2015;208(2):507–518.

10. van der Does HC, Lievens B, Claes L, Houterman PM, Cornelissen BJC, Rep M. The presence of a virulence locus discriminates *Fusarium oxysporum* isolates causing tomato wilt from other isolates. Environ Microbiol. 2008;10(6):1475–1485.

11. Gawehns F, Ma L, Bruning O, Houterman PM, Boeren S, Cornelissen BJC, et al. The effector repertoire of *Fusarium oxysporum* determines the tomato xylem proteome composition following infection. Front Plant Sci. 2015;6:967.

12. Li E, Wang G, Xiao J, Ling J, Yang Y, Xie B. A *SIX1* homolog in *Fusarium oxysporum* f. sp. *conglutinans* is required for full virulence on cabbage. PLoS One. 2016;11(3):e0152273.

13. Thatcher LF, Gardiner DM, Kazan K, Manners JM. A highly conserved effector in *Fusarium oxysporum* is required for full virulence on Arabidopsis. Mol Plant Microbe Interact. 2012;25(2):180–190.

14. An B, Hou X, Guo Y, Zhao S, Luo H, He C, et al. The effector SIX8 is required for virulence of *Fusarium oxysporum* f. sp. *cubense* tropical race 4 to Cavendish banana. Fungal Biol. 2019;123(5):423–430.

15. Widinugraheni S, Nino-Sanchez J, van der Does HC, van Dam P, Garcia-Bastidas FA, Subandiyah S, et al. A SIX1 homolog in *Fusarium oxysporum* f. sp. *cubense* tropical race 4 contributes to virulence towards Cavendish banana. PLoS One. 2018;13(10):e0205896.

16. Cao L, Blekemolen MC, Tintor N, Cornelissen BJC, Takken FLW. The *Fusarium oxysporum* Avr2-Six5 Effector Pair Alters Plasmodesmatal Exclusion Selectivity to Facilitate Cell-to-Cell Movement of Avr2. Mol Plant. 2018;11(5):691–705.

17. Ayukawa Y, Asai S, Gan P, Tsushima A, Ichihashi Y, Shibata A, et al. A pair of effectors encoded on a conditionally dispensable chromosome of *Fusarium oxysporum* suppress host-specific immunity. Commun Biol. 2021;4(1):707.

18. Gonzalez-Cendales Y, Catanzariti AM, Baker B, McGrath DJ, Jones DA. Identification of *I-7* expands the repertoire of genes for resistance to *Fusarium* wilt in tomato to three resistance gene classes. Mol Plant Pathol. 2016;17(3):448–463.

19. Catanzariti AM, Do HTT, Bru P, de Sain M, Thatcher LF, Rep M, et al. The tomato *I* gene for *Fusarium* wilt resistance encodes an atypical leucine-rich repeat receptor-like protein whose function is nevertheless dependent on *SOBIR1* and *SERK3/BAK1*. Plant J. 2017;89(6):1195–1209.

20. Simons G, Groenendijk J, Wijbrandi J, Reijans M, Groenen J, Diergaarde P, et al. Dissection of the *Fusarium I-2* gene cluster in tomato reveals six homologs and one active gene copy. Plant Cell. 1998;10(6):1055–1068.

21. Catanzariti AM, Lim GTT, Jones DA. The tomato *I-3* gene: a novel gene for resistance to *Fusarium* wilt disease. New Phytol. 2015;207(1):106–118.

22. Houterman PM, Cornelissen BJC, Rep M. Suppression of plant resistance gene-based immunity by a fungal effector. PLoS Pathog. 2008;4(5):e1000061.

23. Houterman PM, Ma L, van Ooijen G, de Vroomen MJ, Cornelissen BJC, Takken FLW, et al. The effector protein Avr2 of the xylem-colonizing fungus *Fusarium oxysporum* activates the tomato resistance protein I-2 intracellularly. Plant J. 2009;58(6):970–978.

24. Di X, Cao L, Hughes RK, Tintor N, Banfield MJ, Takken FLW. Structure-function analysis of the *Fusarium oxysporum* Avr2 effector allows uncoupling of its immune-suppressing activity from recognition. New Phytol. 2017;216(3):897–914.

25. Sarma GN, Manning VA, Ciuffetti LM, Karplus PA. Structure of Ptr ToxA: an RGD-containing host-selective toxin from *Pyrenophora tritici-repentis*. Plant Cell. 2005;17(11):3190–3202.

26. Wang CI, Guncar G, Forwood JK, Teh T, Catanzariti AM, Lawrence GJ, et al. Crystal structures of flax rust avirulence proteins AvrL567-A and -D reveal details of the structural basis for flax disease resistance specificity. Plant Cell. 2007;19(9):2898–2912.

27. de Guillen K, Ortiz-Vallejo D, Gracy J, Fournier E, Kroj T, Padilla A. Structure analysis uncovers a highly diverse but structurally conserved effector family in phytopathogenic fungi. PLoS Pathog. 2015;11(10):e1005228.

28. Spanu PD. Cereal immunity against powdery mildews targets RNase-Like Proteins associated with Haustoria (RALPH) effectors evolved from a common ancestral gene. New Phytol. 2017;213(3):969–971.

29. Lazar N, Mesarich CH, Petit-Houdenot Y, Talbi N, Li de la Sierra-Gallay I, Zelie E, et al. A new family of structurally conserved fungal effectors displays epistatic interactions with plant resistance proteins. PLoS Pathog. 2022;18(7):e1010664.

30. Outram MA, Sung YC, Yu D, Dagvadorj B, Rima SA, Jones DA, et al. The crystal structure of SnTox3 from the necrotrophic fungus *Parastagonospora nodorum* reveals a unique effector fold and provides insight into Snn3 recognition and pro-domain protease processing of fungal effectors. New Phytol. 2021;231(6):2282–2296.

31. Outram MA, Solomon PS, Williams SJ. Pro-domain processing of fungal effector proteins from plant pathogens. PLoS Pathog. 2021;17(10):e1010000.

32. Yu D, Outram MA, Creen E, Smith A, Sung YC, Darma R, et al. Optimised production of disulfide-bonded fungal effectors in E. coli using CyDisCo and FunCyDisCo co-expression approaches. Mol Plant Microbe Interact. 2021.

33. Holm L. Dali server: structural unification of protein families. Nucleic Acids Research. 2022;50(W1):W210–W215.

34. Jumper J, Evans R, Pritzel A, Green T, Figurnov M, Ronneberger O, et al. Highly accurate protein structure prediction with AlphaFold. Nature. 2021;596:583–589.

35. Batson AM, Fokkens L, Rep M, du Toit LJ. Putative Effector Genes Distinguish Two Pathogenicity Groups of Fusarium oxysporum f. sp. spinaciae. Mol Plant Microbe Interact. 2021;34(2):141–156.

36. Czislowski E, Fraser-Smith S, Zander M, O’Neill WT, Meldrum RA, Tran-Nguyen LTT, et al. Investigation of the diversity of effector genes in the banana pathogen, Fusarium oxysporum f. sp. cubense, reveals evidence of horizontal gene transfer. Mol Plant Pathol. 2018;19(5):1155–1171.

37. Lievens B, Houterman PM, Rep M. Effector gene screening allows unambiguous identification of Fusarium oxysporum f. sp. lycopersici races and discrimination from other formae speciales. FEMS Microbiol Lett. 2009;300(2):201–215.

38. van Dam P, Fokkens L, Schmidt SM, Linmans JH, Kistler HC, Ma LJ, et al. Effector profiles distinguish formae speciales of Fusarium oxysporum. Environ Microbiol. 2016;18(11):4087–4102.

39. Rocafort M, Bowen JK, Hassing B, Cox MP, McGreal B, de la Rosa S, et al. The *Venturia inaequalis* effector repertoire is dominated by expanded families with predicted structural similarity, but unrelated sequence, to avirulence proteins from other plant-pathogenic fungi. BMC Biology. 2022;20(1):246.

40. Varadi M, Anyango S, Deshpande M, Nair S, Natassia C, Yordanova G, et al. AlphaFold Protein Structure Database: massively expanding the structural coverage of protein-sequence space with high-accuracy models. Nucleic Acids Res. 2022;50(D1):D439–D444.

41. van Kempen M, Kim SS, Tumescheit C, Mirdita M, Lee J, Gilchrist CLM, et al. Fast and accurate protein structure search with Foldseek. Nature Biotechnology. 2023.

42. Iwaoka R, Nagata T, Tsuda K, Imai T, Okano H, Kobayashi N, et al. Structural insight into the recognition of r(UAG) by Musashi-1 RBD2, and construction of a model of Musashi-1 RBD1-2 bound to the minimum target RNA. Molecules. 2017;22(7):1207.

43. Allen A, Chatt E, Smith TJ. The atomic structure of the virally encoded antifungal protein, KP6. J Mol Biol. 2013;425(3):609–621.

44. Li J, Fokkens L, Conneely LJ, Rep M. Partial pathogenicity chromosomes in *Fusarium oxysporum* are sufficient to cause disease and can be horizontally transferred. Environ Microbiol. 2020;22(12):4985–5004.

45. Sun X, Fang X, Wang D, Jones DA, Ma L. Transcriptome Analysis of Fusarium-Tomato Interaction Based on an Updated Genome Annotation of Fusarium oxysporum f. sp. lycopersici Identifies Novel Effector Candidates That Suppress or Induce Cell Death in Nicotiana benthamiana. J Fungi (Basel). 2022;8(7).

46. Evans R, O’Neill M, Pritzel A, Antropova N, Senior A, Green T, et al. Protein complex prediction with AlphaFold-Multimer. bioRxiv. 2022:2021.2010.2004.463034.

47. Mirdita M, Schutze K, Moriwaki Y, Heo L, Ovchinnikov S, Steinegger M. ColabFold: making protein folding accessible to all. Nat Methods. 2022;19(6):679–682.

48. Stergiopoulos I, de Wit PJGM. Fungal effector proteins. Annu Rev Phytopathol. 2009;47:233–263.

49. Outram MA, Figueroa M, Sperschneider J, Williams SJ, Dodds PN. Seeing is believing: Exploiting advances in structural biology to understand and engineer plant immunity. Curr Opin Plant Biol. 2022;67:102210.

50. Pennington HG, Jones R, Kwon S, Bonciani G, Thieron H, Chandler T, et al. The fungal ribonuclease-like effector protein CSEP0064/BEC1054 represses plant immunity and interferes with degradation of host ribosomal RNA. PLoS Pathog. 2019;15(3):e1007620.

51. Pedersen C, Ver Loren van Themaat E, McGuffin LJ, Abbott JC, Burgis TA, Barton G, et al. Structure and evolution of barley powdery mildew effector candidates. BMC Genomics. 2012;13:694.

52. Blondeau K, Blaise F, Graille M, Kale SD, Linglin J, Ollivier B, et al. Crystal structure of the effector AvrLm4-7 of *Leptosphaeria maculans* reveals insights into its translocation into plant cells and recognition by resistance proteins. Plant J. 2015;83(4):610–624.

53. Petit-Houdenot Y, Langner T, Harant A, Win J, Kamoun S. A clone resource of *Magnaporthe oryzae* effectors that share sequence and structural similarities across host-specific lineages. Mol Plant Microbe Interact. 2020;33(8):1032–1035.

54. Teulet A, Quan C, Evangelisti E, Wanke A, Yang W, Schornack S. A pathogen effector FOLD diversified in symbiotic fungi. New Phytol. 2023;239(3):1127–1139.

55. Bauer S, Yu D, Lawson AW, Saur IML, Frantzeskakis L, Kracher B, et al. The leucine-rich repeats in allelic barley MLA immune receptors define specificity towards sequence-unrelated powdery mildew avirulence effectors with a predicted common RNase-like fold. PLoS Pathog. 2021;17(2):e1009223.

56. Seong K, Krasileva K. Computational structural genomics unravels common folds and novel families in the secretome of fungal phytopathogen *Magnaporthe oryzae*. Mol Plant Microbe Interact. 2021.

57. Yan X, Tang B, Ryder LS, MacLean D, Were VM, Eseola AB, et al. The transcriptional landscape of plant infection by the rice blast fungus Magnaporthe oryzae reveals distinct families of temporally co-regulated and structurally conserved effectors. Plant Cell. 2023;35(5):1360–1385.

58. Seong K, Krasileva KV. Prediction of effector protein structures from fungal phytopathogens enables evolutionary analyses. Nat Microbiol. 2023;8(1):174–187.

59. Ortiz D, Chen J, Outram MA, Saur IML, Upadhyaya NM, Mago R, et al. The stem rust effector protein AvrSr50 escapes Sr50 recognition by a substitution in a single surface-exposed residue. New Phytol. 2022;234(2):592–606.

60. Park SY, Jeong MS, Park SA, Ha SC, Na BK, Jang SB. Structural basis of the cystein protease inhibitor *Clonorchis sinensis* Stefin-1. Biochem Biophys Res Commun. 2018;498(1):9–17.

61. Renko M, Taler-Vercic A, Mihelic M, Zerovnik E, Turk D. Partial rotational lattice order-disorder in stefin B crystals. Acta Crystallogr D Biol Crystallogr. 2014;70(Pt 4):1015–1025.

62. Dietrich JD, Longenecker KL, Wilson NS, Goess C, Panchal SC, Swann SL, et al. Development of orally efficacious allosteric inhibitors of TNFalpha via fragment-based drug design. J Med Chem. 2021;64(1):417–429.

63. Sulak O, Cioci G, Delia M, Lahmann M, Varrot A, Imberty A, et al. A TNF-like trimeric lectin domain from *Burkholderia cenocepacia* with specificity for fucosylated human histo-blood group antigens. Structure. 2010;18(1):59–72.

64. Khoshnevis S, Neumann P, Ficner R. Crystal structure of the RNA recognition motif of yeast translation initiation factor eIF3b reveals differences to human eIF3b. PLoS One. 2010;5(9):e12784.

65. Bull PC, Cox DW. Wilson disease and Menkes disease: new handles on heavy-metal transport. Trends Genet. 1994;10(7):246–252.

66. Bleackley MR, Vasa S, Harvey PJ, Shafee TMA, Kerenga BK, Soares da Costa TP, et al. Histidine-rich defensins from the *Solanaceae* and *Brasicaceae* are antifungal and metal binding proteins. J Fungi 2020;6(3):145.

67. Lay FT, Ryan GF, Caria S, Phan TK, Veneer PK, White JA, et al. Structural and functional characterization of the membrane-permeabilizing activity of *Nicotiana occidentalis* defensin NoD173 and protein engineering to enhance oncolysis. FASEB J. 2019;33(5):6470–6482.

68. Prochnicka-Chalufour A, Corzo G, Satake H, Martin-Eauclaire MF, Murgia AR, Prestipino G, et al. Solution structure of discrepin, a new K+-channel blocking peptide from the alpha-KTx15 subfamily. Biochemistry. 2006;45(6):1795–1804.

69. Korolkova YV, Bocharov EV, Angelo K, Maslennikov IV, Grinenko OV, Lipkin AV, et al. New binding site on common molecular scaffold provides HERG channel specificity of scorpion toxin BeKm-1. J Biol Chem. 2002;277(45):43104–43109.

70. Van Duyne GD, Ghosh G, Maas WK, Sigler PB. Structure of the oligomerization and L-arginine binding domain of the arginine repressor of *Escherichia coli*. J Mol Biol. 1996;256(2):377–391.

71. Cherney LT, Cherney MM, Garen CR, Lu GJ, James MN. Structure of the C-terminal domain of the arginine repressor protein from *Mycobacterium tuberculosis*. Acta Crystallogr D Biol Crystallogr. 2008;64(Pt 9):950–956.

72. Forderer A, Li E, Lawson AW, Deng YN, Sun Y, Logemann E, et al. A wheat resistosome defines common principles of immune receptor channels. Nature. 2022;610(7932):532-539.

73. Ma S, Lapin D, Liu L, Sun Y, Song W, Zhang X, et al. Direct pathogen-induced assembly of an NLR immune receptor complex to form a holoenzyme. Science. 2020;370(6521):eabe3069.

74. Martin R, Qi T, Zhang H, Liu F, King M, Toth C, et al. Structure of the activated ROQ1 resistosome directly recognizing the pathogen effector XopQ. Science. 2020;370(6521):eabd9993.

75. Plissonneau C, Daverdin G, Ollivier B, Blaise F, Degrave A, Fudal I, et al. A game of hide and seek between avirulence genes *AvrLm4-7* and *AvrLm3* in *Leptosphaeria maculans*. New Phytol. 2016;209(4):1613–1624.

76. Ghanbarnia K, Ma L, Larkan NJ, Haddadi P, Fernando WGD, Borhan MH. *Leptosphaeria maculans* AvrLm9: a new player in the game of hide and seek with AvrLm4-7. Mol Plant Pathol. 2018;19(7):1754–1764.

77. Haddadi P, Larkan NJ, Van deWouw A, Zhang Y, Xiang Neik T, Beynon E, et al. *Brassica napus* genes *Rlm4* and *Rlm7*, conferring resistance to *Leptosphaeria maculans*, are alleles of the *Rlm9* wall-associated kinase-like resistance locus. Plant Biotechnol J. 2022;20(7):1229–1231.

78. Larkan NJ, Ma L, Haddadi P, Buchwaldt M, Parkin IAP, Djavaheri M, et al. The *Brassica napus* wall-associated kinase-like (WAKL) gene *Rlm9* provides race-specific blackleg resistance. Plant J. 2020;104(4):892–900.

79. Bentham AR, Youles M, Mendel MN, Varden FA, De la Concepcion JCB, Mark J. pOPIN-GG: A resource for modular assembly in protein expression vectors. bioRxiv. 2021:2021.2008.2010.455798.

80. Iverson SV, Haddock TL, Beal J, Densmore DM. CIDAR MoClo: improved MoClo assembly standard and new *E. coli* part library enable rapid combinatorial design for synthetic and traditional biology. ACS Synth Biol. 2016;5(1):99–103.

81. Wiedemann C, Bellstedt P, Gorlach M. CAPITO--a web server-based analysis and plotting tool for circular dichroism data. Bioinformatics. 2013;29(14):1750–1757.

82. Cowieson NP, Aragao D, Clift M, Ericsson DJ, Gee C, Harrop SJ, et al. MX1: a bending-magnet crystallography beamline serving both chemical and macromolecular crystallography communities at the Australian Synchrotron. J Synchrotron Radiat. 2015;22(1):187–190.

83. Kabsch W. XDS. Acta Crystallogr D Biol Crystallogr. 2010;66(Pt 2):125–132.

84. Evans PR, Murshudov GN. How good are my data and what is the resolution? Acta Crystallogr D Biol Crystallogr. 2013;69(Pt 7):1204–1214.

85. Winn MD, Ballard CC, Cowtan KD, Dodson EJ, Emsley P, Evans PR, et al. Overview of the *CCP4* suite and current developments. Acta Crystallogr D Biol Crystallogr. 2011;67(Pt 4):235–242.

86. Skubak P, Pannu NS. Automatic protein structure solution from weak X-ray data. Nat Commun. 2013;4:2777.

87. Skubak P, Arac D, Bowler MW, Correia AR, Hoelz A, Larsen S, et al. A new MR-SAD algorithm for the automatic building of protein models from low-resolution X-ray data and a poor starting model. IUCrJ. 2018;5(Pt 2):166–171.

88. Afonine PV, Grosse-Kunstleve RW, Echols N, Headd JJ, Moriarty NW, Mustyakimov M, et al. Towards automated crystallographic structure refinement with *phenix.refine*. Acta Crystallogr D Biol Crystallogr. 2012;68(Pt 4):352–367.

89. Emsley P, Lohkamp B, Scott WG, Cowtan K. Features and development of *COOT*. Acta Crystallogr D Biol Crystallogr. 2010;66(Pt 4):486–501.

90. Aragao D, Aishima J, Cherukuvada H, Clarken R, Clift M, Cowieson NP, et al. MX2: a high-flux undulator microfocus beamline serving both the chemical and macromolecular crystallography communities at the Australian Synchrotron. J Synchrotron Radiat. 2018;25(Pt 3):885–891.

91. Terwilliger TC, Grosse-Kunstleve RW, Afonine PV, Moriarty NW, Zwart PH, Hung LW, et al. Iterative model building, structure refinement and density modification with the *PHENIX AutoBuild* wizard. Acta Crystallogr D Biol Crystallogr. 2008;64(Pt 1):61–69.

92. Almagro Armenteros JJ, Tsirigos KD, Sonderby CK, Petersen TN, Winther O, Brunak S, et al. SignalP 5.0 improves signal peptide predictions using deep neural networks. Nat Biotechnol. 2019;37(4):420–423.

93. Hellens RP, Edwards EA, Leyland NR, Bean S, Mullineaux PM. pGreen: a versatile and flexible binary Ti vector for *Agrobacterium*-mediated plant transformation. Plant Mol Biol. 2000;42(6):819–832.

94. Stivala A, Wybrow M, Wirth A, Whisstock JC, Stuckey PJ. Automatic generation of protein structure cartoons with Pro-origami. Bioinformatics. 2011;27(23):3315–3316.

