## Supplementary methods for "The structural repertoire of *Fusarium oxysporum* f. sp. *lycopersici* effectors revealed by experimental and computational studies"

**Crystallisation screening and optimisation**

Initial screening to determine crystallisation conditions was performed at a concentration of 9.5 mg/mL for Avr3^22-284^, 10 mg/mL for Avr1^18-242^, Avr1^59-242^, SIX8^50-141^ and PSL1^18-111^, 15 mg/mL for SIX6^17-225^ and SIX6^58-225^, 25 mg/mL for SIX8_C58S^19-141^, 18 mg/mL for SIX8_C58S^50-141^ and PSL1_C37S^18-111^, 14 mg/mL for SIX8-PSL1 complex and SIX13 with and without Kex2 protease in 96-well MRC 2 plates (Hampton Research) at 18°C using the sitting-drop vapour-diffusion method and commercially available sparse matrix screens. For screening, 150 nL protein solution and 150 nL reservoir solution was prepared on a sitting-drop well using an NT8®-Drop Setter robot (Formulatrix, USA). The drops were monitored and imaged using the Rock Imager system (Formulatrix, USA) over the course of a month.

For Avr1^18-242^, SIX6^17-225^, SIX8^50-141^, PSL1^18-111^, SIX8-PSL1 complex and SIX13^22-293^, no crystals were obtained from the different sparse matrix screens trialled. From initial screening, crystals with the best morphology for Avr3^22-284^ were obtained in (1) 0.2 M lithium sulfate, 0.1 M Bis-Tris pH 6.5 and 25% (w/v) PEG 3350 (SG1 screen: condition D10), and (2) 0.2 M ammonium sulfate, 0.1 M Bis-Tris pH 6.5 and 25% (w/v) PEG 3350 (SG1 screen: condition F5). Crystals were visible after a period of 3 days and continued to grow for 3 weeks after initial setup. Replicate drops with 1 μl protein solution at 9.5 mg/mL and 1 μl reservoir solution were set-up in 24-well hanging-drop vapour-diffusion plates and produced crystals within 4 days that continued to grow over 1 month. No crystal optimisation was needed for Avr3, with the final conditions being (1) 0.2 M ammonium sulfate, 0.1 M Bis-Tris pH 6.5, 25% (w/v) PEG 3350, and (2) 0.2 M lithium sulfate, 0.1 M Bis-Tris pH 6.5, 25% (w/v) PEG 3350. For Avr1^59-242^, crystals with the best morphology were obtained in (1) 0.2 M ammonium sulfate, 0.1 M sodium acetate pH 4.6, 25% (w/v) PEG 4000 (SG1 screen: condition C1) and (2) 0.2 M ammonium sulfate, 30% (w/v) PEG 8000 (SG1 screen: condition D7) within 1 day of initial setup. Crystal optimisation was carried out in 24-well hanging-drop vapour-diffusion plates at 18°C. The final optimised condition for Avr1^59-242^ was 0.2 M ammonium sulfate, 0.1 M sodium acetate pH 4.5, 17.5% (w/v) PEG 4000 at a protein concentration of 7 mg/mL with microseeding over a period of 3 weeks. For SIX6^58-225^, crystals were obtained in 0.2 M ammonium tartrate and 20% (w/v) PEG 3350 (SG1 screen: condition G9) 40 days after initial setup. Crystals were picked directly from the sparse matrix screen. For SIX8_C58S^50-141^, crystals were obtained in 0.17 M ammonium sulfate, 15% (v/v) glycerol and 25.5% (w/v) PEG 4000 (JCSG screen: condition D9) a week after initial setup. Crystals were picked directly from the sparse matrix screen. For SIX13, Kex2 protease was added to the protein at a 1:200 protease to protein ratio prior to crystal tray setup. Crystals with the best morphology were obtained in (1) 0.2 M lithium sulfate, 0.1 M Bis-Tris pH 6.5 and 25% (w/v) PEG 3350 (SG1 screen: condition D10), and (2) 0.2 M ammonium sulfate, 0.1 M Bis-Tris pH 6.5 and 25% (w/v) PEG 3350 (SG1 screen: condition F5) within 2 days of initial setup. Crystals were optimised using hanging-drop vapour-diffusion plates and the final optimised condition for SIX13 was 0.2 M lithium sulfate, 0.1 M Bis-Tris pH 6.5, 25% (w/v) PEG 3350 at a protein concentration of 14 mg/mL. For PSL1_C37S^18-111^, crystals were obtained in 70% (v/v) MPD and 0.1 M HEPES pH 7.5 within 3 days after initial setup. Crystal optimisation was carried out in 24-well hanging-drop vapour-diffusion plates at 18°C. The final optimised condition for PSL1_C37S^18-111^ was 62% (v/v) MPD and 0.1 M HEPES pH 7.5 at a protein concentration of 17.5 mg/mL.
